# Spontaneously Arising *Streptococcus mutans* Variants With Reduced Susceptibility to Chlorhexidine Display Genetic Defects and Diminished Fitness

**DOI:** 10.1101/528471

**Authors:** Justin R. Kaspar, Matthew J. Godwin, Irina M. Velsko, Vincent P. Richards, Robert A. Burne

**Author notes:** Corresponding author Mailing address: Department of Oral Biology, University of Florida, College of Dentistry, P.O. Box 100424, Gainesville, FL 32610.

## Abstract

Chlorhexidine (CHX) has been used to control dental caries, caused by acid-tolerant bacteria such as *Streptococcus mutans*, since the 1970s. Repeated CHX exposures in other bacterial species results in development of variants with reduced susceptibility that also become more resistant to other antimicrobials. It has not been tested if such variants arise when streptococci are exposed to CHX. Here, we passage *S. mutans* in increasing concentrations of CHX and isolate spontaneously-arising <u>r</u>educed <u>s</u>usceptibility <u>v</u>ariants (RSVs) from separate lineages that have minimal inhibitory concentrations (MICs) that are up to 3-fold greater than the parental strain. The RSVs have increased growth rates at neutral pH and under acidic conditions in the presence of CHX, but accumulate less biomass in biofilms. RSVs display higher MICs for daptomycin and clindamycin, but increased sensitivity to dental-relevant antimicrobials triclosan and sodium fluoride. Plate-based assays for competition with health-associated oral streptococci reveal decreased bacteriocin production by the RSVs, increased sensitivity to hydrogen peroxide, and diminished competitive fitness in a human-derived *ex vivo* biofilm consortium. Whole genome sequencing identified common single nucleotide polymorphisms (SNPs) within a diacylglycerol kinase homolog and a glycolipid synthesis enzyme, which could alter accumulation of lipoteichoic acids and other envelope constituents; as well as a variety of mutations in other genes. Collectively, these findings confirm that *S. mutans* and likely other streptococci can develop tolerance to CHX, but that increased tolerance comes at a fitness cost, such that CHX-induced variants that spontaneously arise in the human oral cavity may not persist.

## INTRODUCTION

Controlling microbial biofilm infections in humans is challenging in part due to the presence of extracellular polymeric substances (EPS) in which the organisms are embedded, the presence of persister cells as well as subpopulations of cells with low metabolic activity, and exchange of antimicrobial resistance genes through horizontal gene transfer. In the human oral cavity, accumulation of biofilms on the teeth can lead to destruction of tooth mineral when there is frequent and significant localized acidification by organic acids produced as metabolic end products from fermentable carbohydrates (1). The initiation and progression of dental caries can be controlled to a degree through non-specific mechanical removal of the biofilms or by treatment with various chemical agents that include fluoride, xylitol or chlorhexidine (CHX) (2, 3). While the effectiveness of CHX in caries prevention is not definitively established and there can be undesirable side effects (e.g. staining, irritation of host tissues) (4), CHX remains a standard against which antimicrobial therapies are compared for their efficacy in the suppression of caries-causing bacteria, such as *Streptococcus mutans* (5–7). Currently in the United States, mouth rinses containing 0.12% CHX are the only CHX-containing products marketed for caries prevention (4).

CHX is a symmetric bis-biguanide agent comprising two chloroguanide chains that are connected by a central hexamethylene chain and has diverse medical applications as a disinfectant for surfaces and as an antiseptic for topical applications. CHX carries two positive charges at physiological pH and is attracted to the bacterial cell surface, where it may electrostatically interact with negatively charged phospholipids (8). Depending on the concentration of CHX, it can reduce bacterial membrane fluidity or disrupt the structural integrity of the membrane(s), causing increased permeability, leakage of cell contents, loss of proton motor force and cell death as its main mode of action (9). In one study (10), it was found that CHX induces the formation of dented spots on the surface of the cell wall of both *Escherichia coli* and *Bacillus subtilis*. The authors posited that the structural changes of the bacterial cell wall may be due to the action of CHX with membrane cardiolipin and phosphatidylethanolamine to disturb the normal arrangement and integrity of the phospholipid bilayer structure and its associated proteins (10). A similar finding was recently made in *S. mutans*, where vesicle-like structures were apparent on the cell surface after treatment with 0.2% CHX (11). In the same study, treatment of 72-hour *S. mutans* biofilms resulted in 80% of the population displaying loss of membrane integrity, assessed by staining with propidium iodide (PI) and analysis of the population using flow cytometry (11).

While CHX likely puts substantial selective pressure on *S. mutans* and the oral microbiota in general, particularly if used repeatedly for oral care, the organisms appear to remain susceptible to CHX, with few reports of resistance (12). However one recent study (6) reported that when CHX was continuously flowed on biofilms formed on polystyrene blocks *in vitro*, detached cells displayed a higher minimal inhibitory concentration (MIC) to CHX (1.25 μg mL^-1^), whereas most host-derived isolates of *S. mutans* evaluated had MICs < 1μg mL^1^ (6). Earlier studies with other oral streptococci, including *Streptococcus sanguinis* and *Streptococcus mitis*, showed that repeated exposure to CHX resulted in increased MICs to CHX, but these findings were not deemed to be of clinical significance (13–17). In enterococci, increased tolerance to antiseptics like CHX in hospital settings has inspired research into mechanisms for increased CHX tolerance, including studies of CHX effects on the transcriptome (18) and analysis of the impact of CHX on a transposon mutant library (19). More specifically, Bhardwaj and colleagues (18) used RNA-Seq to show that VanA-type vancomycin resistance genes are upregulated in *Enterococcus faecium* following exposure to CHX. In *E. faecium*, two genes encoding for a two-component regulatory system were found to contribute to CHX tolerance (19). Thus, it is clear that Firmicutes have the capacity to develop increased tolerance to CHX, but the mechanisms by which tolerance may develop, especially in streptococci, is only beginning to be explored.

There is a continued need to be alert to the emergence of new oral isolates that display reduced susceptibility to CHX due to its increased usage within the clinic (20). Adaptations to CHX may also lead to cross-resistance to other antimicrobials, as was noted recently with clinical strains of *Klebsiella pneumoniae* that could acquire resistance to the last-resort antibiotic colistin after exposure to CHX (21). To date, the emergence of tolerant or resitant oral bacteria due to repeated use of CHX in the dental practice has not been fully studied, although determination of whether exposure to CHX can result in acquisition of resistance to other antimicrobials and the molecular mechanisms by which CHX may confer resistance in these organisms is of growing interest (22). In this study, we focus on the caries-associated pathogen *S. mutans*, show that passage on increasing concentrations of CHX gives rise to variants that display reduced susceptibility to CHX, and begin to examine the genetic basis for reduced CHX susceptibility. This research serves as a starting point to investigate the mechanisms of action of CHX and how streptococci cope with this chemical stressor that will allow us to generate information that supports the development of more effective treatments for removal of cariogenic bacteria from oral biofilm communities.

## RESULTS AND DISCUSSION

### Selection for Chlorhexidine Reduced Susceptibility Variants

To begin our study, we initially screened nine strains of *S. mutans* that were acquired from different sources and that displayed phenotypic heterogeneity in established virulence-related traits (23, 24). The MICs for the nine strains ranged from 0.25 – 0.625 μg mL^-1^ CHX **(Supplemental Figure 1)**, and none greatly exceeded the previously reported 0.5 μg mL^-1^ MIC for *S. mutans* (12). For the continuation of the research, we chose to work with strain UA159, which was orginally isolated in 1982 from a child with active caries and serves as the wild-type strain for many laboratories that study *S. mutans*. We next wanted to determine how *S. mutans* may adapt to repeated CHX exposures over time, and whether isolates that display reduced susceptibility may arise. To accomplish this, UA159 was passaged on BHI agar plates containing various concentrations of CHX **(Figure 1A)**. Colonies able to grow on the highest concentration CHX agar plate were picked, regrown in BHI broth under non-selective conditions, and then plated on a higher concentration of CHX. The selections were then repeated with one additional round using higher concentrations of CHX and the colonies were stored for further analysis.

**Figure 1.**
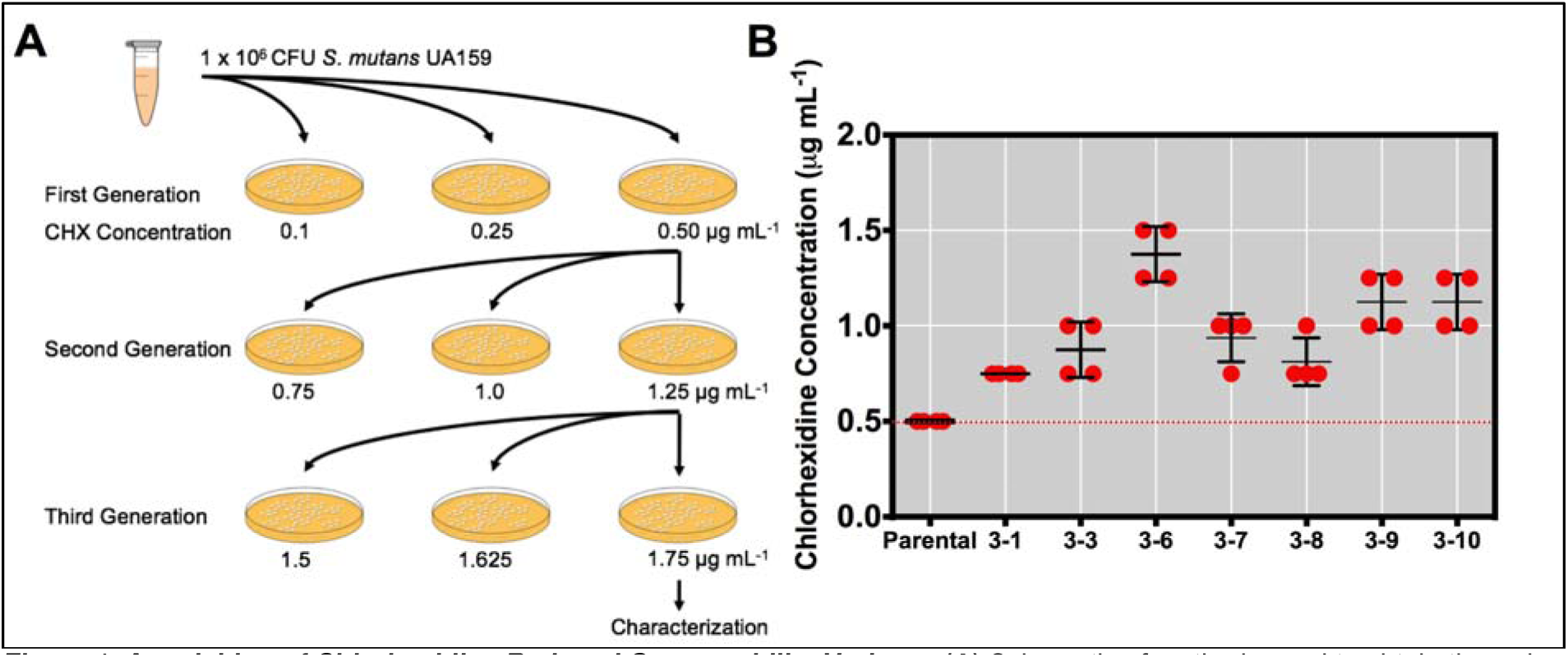
Acquisition of Chlorhexidine Reduced Susceptability Variants. **(A)** Schematic of methods used to obtain the reduced susceptibility variants to chlorhexidine (RSVs). ∼1×10^6^ CFU of *S. mutans* UA159 grown to mid-exponential phase was spread plated on BHI agar plates containing either 0.1, 0.25 or 0.5 μg mL^-1^ CHX. Ten different single colonies were selected from the 0.5 μg mL^-1^ CHX plate to establish 10 independent lineages (RSV1). Cultures of these colonies were then plated on separate 0.75, 1.0 or 1.25 μg mL^-1^ CHX BHI agar plates (RSV2). One randomly selected colony from each plate was then cultured in BHI and plated on separate 1.5, 1.625 or 1.75 μg mL^-1^ CHX. At this point, one randomly selected colony from each of the RSV3 plates was stored for further analysis and characterization. **(B)** MIC determination of the parental along with the RSV3s grown for 24 h in BHI with the designated concentrations of CHX. Red dots denote highest CHX concentration that the strains displayed growth (OD_600 nm_ > 0.1). Four biological replicates of each isolate were tested during the experiment and was replicated four times.

To acquire the first generation of spontaneously-arising CHX reduced <u>s</u>usceptibility *S. mutans* <u>v</u>ariants (RSVs), 1×10^6^ colonly forming units (CFUs) of *S. mutans* UA159 grown in BHI broth to OD_600 nm_ = 0.5 (mid-exponential log phase) was spread on BHI agar plates containing 0.1, 0.25 or 0.5 μg mL^-1^ CHX. While the plates containing 0.1 or 0.25 μg mL^-1^ CHX yielded lawns of bacteria, individual colonies were obtained on the 0.5 μg mL^-1^ agar plates. No colonies could be obtained in the first round of plating on BHI agar plates supplemented with 0.75 μg mL^-1^ CHX. Ten individual bacterial colonies were selected from the 0.5 μg mL^-1^ plate and designated as RSV1s (reduced susceptibility variants from the 1^st^ generation). These colonies were regrown to OD_600 nm_ = 0.5 in broth, diluted to 1×10^6^ CFUs and 0.1 mL of each of the 10 RSV1 cultures were spread on individual BHI agar plates containing either 0.75, 1.0 or 1.25 μg mL^-1^ CHX, establishing 10 independent lineages moving forward to the second generation (RSV2s). Individual colonies were obtained on plates with all three concentrations of CHX. One colony was randomly selected from each of the 1.25 μg mL^-1^ CHX plates to move forward to the third generation (RSV3s). After growth to OD_600 nm_ = 0.5 and dilution to 1×10^6^ CFUs, these 10 RSV2s were then plated on 1.5, 1.625 or 1.75 μg mL^-1^ CHX. Again, colonies were obtained on all concentrations. At this point, one randomly-selected colony from each of the RSV3 plates (1.75 μg mL^-1^ CHX) was stored for further analysis. Confirmation that the variants were indeed *S. mutans* was obtained by amplifying and sequencing a 1.5 kbp region from the 16S rDNA sequence. Seven selected RSV3s that returned as *S. mutans* were subjected to further analysis.

To confirm that the selected RSV3s had reduced susceptibility to CHX, an MIC test was conducted **(Figure 1B)**. While the parental strain could only grow in BHI broth containing 0.5 μg mL^-1^ CHX after 24 h, all RSV3s were able to grow in BHI containing >0.5 μg mL^-1^ CHX. Interestingly, all seven RSV3s tested displayed variable MICs, with the most tolerant isolate, RSV3-6, able to grow in BHI broth containing 1.50 μg mL^-1^ CHX, a 3-fold increase from the parental strain. Other isolates, such as RSV3-1 and RSV3-8 were only able to grow in BHI broth containing 0.75 μg mL^-1^ CHX after 24 h, a half-fold increase from parental. These data indicated to us that the seven selected RSV3s are diverse in that they were able to propagate in different concentrations of CHX, possibly indicative of the acquisition of different mutations within the genome.

### Attributes of the RSV3s

We reasoned that potential mutations could have pleiotropic effects, so several key properties of the variants were examined. We found that a majority of the RSV3s, except RSV3-1 and RSV3-9, had a shorter lag phase compared to the parental UA159 **(Figure 2A)**. However, all RSV3s displayed longer doubling times than the parental strain, ranging from 56 ± 2 min for RSV3-3 to 145 ± 26 min for RSV3-1, in contrast to 51 ± 2 for the parental **(Table 1)**. When growth rates in the presence of 0.75 μg mL^-1^ CHX were determined, the parental strain displayed little growth over a 24 h period (doubling time 623 ± 69 min, final yield 0.29 ± 0.09 OD_600 nm_), whereas all isolates displayed average doubling times under 150 min and final yields higher than 0.45 **(Figure 2B)**. Some variants, including RSV3-3, RSV3-8 and RSV 3-10, showed little difference in doubling times in the presence or absence of 0.75 μg mL^-1^ CHX. These data confirm that the selected isolates have reduced susceptibility to CHX, and provide support for the idea that the disparate growth characteristics may reflect variation in the underlying genetic and physiologic bases for increased MICs to CHX.

**TABLE 1.** Growth characteristics of RSV3s under different conditions.

| Strain | Growth Characteristics |  |  |  |  |  |  |  |
| --- | --- | --- | --- | --- | --- | --- | --- | --- |
|  | BHI pH 7.0 |  | BHI pH 7.0 + 0.75 µg mL<sup>-1</sup> CHX |  | BHI pH 5.5 |  | BHI pH 5.5 + 0.75 µg mL<sup>-1</sup> CHX |  |
|  | Doubling Time (min) | Final Yield (OD<sub>600 nm</sub>) | Doubling Time (min) | Final Yield (OD<sub>600 nm</sub>) | Doubling Time (min) | Final Yield (OD<sub>600 nm</sub>) | Doubling Time (min) | Final Yield (OD<sub>600 nm</sub>) |
| Parental | 51 ± 2 | 0.61 ± 0.08 | 623 ± 69 | 0.29 ± 0.09 | 178 ± 12 | 0.65 ± 0.01 | No growth | 0.04 ± 0.02 |
| RSV3-1 | 145 ± 26 | 0.45 ± 0.11 | 126 ± 19 | 0.41 ± 0.10 | 186 ± 11 | 0.45 ± 0.02 | 142 ± 14 | 0.23 ± 0.07 |
| RSV3-3 | 56 ± 2 | 0.73 ± 0.14 | 78 ± 7 | 0.50 ± 0.04 | 259 ± 25 | 0.41 ± 0.01 | 142 ± 12 | 0.48 ± 0.02 |
| RSV3-6 | 141 ± 16 | 0.55 ± 0.08 | 139 ± 17 | 0.44 ± 0.05 | 132 ± 14 | 0.32 ± 0.02 | 145 ± 8 | 0.31 ± 0.02 |
| RSV3-7 | 108 ± 11 | 0.67 ± 0.05 | 143 ± 32 | 0.57 ± 0.11 | 142 ± 44 | 0.47 ± 0.02 | 119 ± 11 | 0.59 ± 0.09 |
| RSV3-8 | 66 ± 7 | 0.72 ± 0.03 | 92 ± 13 | 0.59 ± 0.11 | 189 ± 34 | 0.54 ± 0.01 | 121 ± 12 | 0.59 ± 0.04 |
| RSV3-9 | 97 ± 8 | 0.63 ± 0.04 | 133 ± 12 | 0.60 ± 0.01 | 160 ± 10 | 0.34 ± 0.01 | 215 ± 23 | 0.39 ± 0.11 |
| RSV3-10 | 63 ± 6 | 0.56 ± 0.12 | 74 ± 5 | 0.52 ± 0.01 | 107 ± 7 | 0.55 ± 0.03 | 113 ± 22 | 0.59 ± 0.03 |

**Figure 2.**
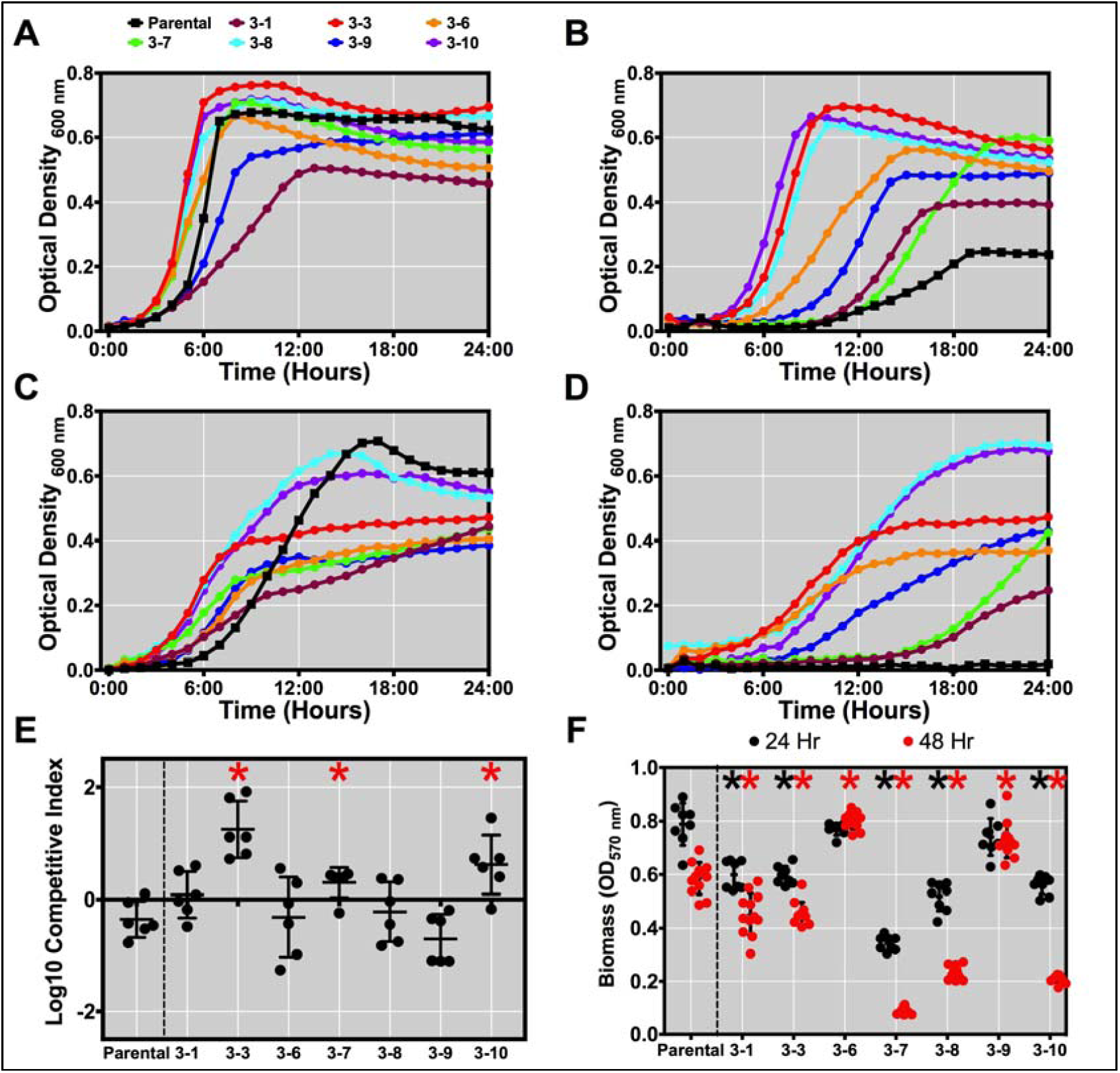
Growth attributes of the RSV3s. Growth of the parental UA159 strain (black line, squares) and each of the RSV3s (colored lines, circles) in either **(A)** BHI, **(B)** BHI with 0.75 μg mL^-1^ CHX, **(C)** BHI pH 5.5, and **(D)** BHI pH 5.5 with 0.75 μg mL^-1^ CHX. **(E)** Competitive index assay between the parental strain with the RSV3s in BHI medium after 24 h, competed with a marked UA159 that included integration of plasmid pMC340B (Km^R^) that was used for selection. Data plotted is in the competitive index (Log10) of the unmarked parental or the RSV3s. * = P value ≤ 0.1 by Student’s T-Test compared to the parental. **(F)** Accumulated biofilm biomass for all individual strains grown for either 24 (black dots) or 48 hours (red dots) in CDM medium supplemented with 20 mM glucose and 5 mM sucrose. 12 individual biofilms from three biological replicates were measured. The experiment was replicated twice.

One critical virulence attribute of *S. mutans* is acid tolerance (aciduricity), which allows organisms to continue metabolic processes at lower pH values than acid-sensitive oral organisms, leading to ecological shifts that promote caries development (25–27). Additionally, when *S. mutans* is cultured at acidic pH values, the fatty acid profile of the membrane is modified from mainly short-chained saturated fatty acids to a membrane with increased proportions of mono-unsaturated fatty acids with longer chains (28). Interestingly, all isolates had a shorter lag time compared to the parental when diluted from BHI at neutral pH into BHI that had been acidified with HCl to a pH of 5.5 **(Figure 2C)**. However, the doubling time of the RSVs was comparable to that of the parental, with the parental obtaining the highest final yield. When 0.75 *µ*g mL^-1^ CHX was included in the acidified broth, the parental strain failed to grow after 48 h, while all variants displayed growth **(Figure 2D)**. Some variants, including RSV3-3 and RSV3-7, actually displayed shorter doubling times and higher final yields when grown with CHX, compared with no CHX present. One possible explanation is that the variants have changes in the the phospholipid profile that allows them to be more toleratant of, and/or more rapidly adapt to, the low pH challenge.

The ability of the RSVs to compete with the parental strain and to form biofilms was also assessed. For competitive index measurements, a marked parental that included integration of an antibiotic resistance marker from pMC340B, which allow for single copy integration of a kanamycin resistance marker in the *mtlA* – *phnA* locus (29), was competed against unmarked parental and the RSV3s **(Figure 2E)**. In planktonic cultures, RSV3-3, RSV3-6 and RSV3-10 were significantly more competitive against the parental strain after 24 h. These differences are consistent with the varying growth rates displayed by the parental and variants. In terms of biofilm formation in the presence of sucrose, all RSV3s other than RSV3-6 and RSV3-9 had significantly less biomass accumulation compared to the parental after 24 h of establishment and growth **(Figure 2F)**. After 48 h, variants RSV3-6 and RSV3-9 showed stable biomass accumulation that now surpassed the parental strain. Interestingly, RSV3-7, RSV3-8 and RSV3-10 displayed a 3-fold decrease in biomass compared to parental strain at the 48 h time point. The mechanisms by which the acquired mutations and/or changes in gene expression impact the biofilm phenotype of the RSV3s will need to be further explored, but the differences observed in biofilm forming capacity futher highlights their diversity.

### RSV3s have reduced susceptibility to other antimicrobials

Two recent studies with *E. faecium* and Methicillin-Resistant *Staphylococcus aureus* (MRSA) demonstrated reduced antibiotic susceptibility in strains that are less sensitive to CHX or the biguanide antiseptic polyhexanide, respectively (30, 31). To determine if reduced susceptibility to CHX could lead to cross-resistance against other antimicrobials in our variants, we determined MICs of the RSV3s for six different antibiotics (amoxicillin, azithromycin, bacitracin, clindamycin, daptomycin, vancomycin) that utilize different modes of action using the Etest Strip Assay **(Table 2)**. We saw no change in susceptibility to antibiotics that inhibit peptidoglycan cross-linking (amoxicillin, vancomycin), but changes against cell-wall targeting bacitracin and daptomycin were evident **(Supplemental Figure 2)**. All variants, except for RSV3-9, displayed increased susceptibility to bacitracin, and all variants except for RSV3-1 showed higher MICs to daptomycin (2-6 fold increase). The latter result is consistent with the fact that CHX and daptomycin disrupt cell membrane potential and fluidity (30, 31). Interestingly, the RSV3s also showed reduced susceptibility to clindamycin, which inhibits protein synthesis through binding to the 50S ribosomal subunit (32), perhaps associated with decreased permeability or increased export of the drug in the RSV3s.

**TABLE 2.** Changes in antibiotic sensitivity for RSV3s.

| Antibiotic | Mechanism of Action | MIC Determined by ETest® Strips (µg mL<sup>-1</sup>) |  |  |  |  |  |  |  |
| --- | --- | --- | --- | --- | --- | --- | --- | --- | --- |
|  |  | Parental | RSV3-1 | RSV3-3 | RSV3-6 | RSV3-7 | RSV3-8 | RSV3-9 | RSV3-10 |
| Amoxicillin | Inhibition of peptidoglycan cross-linkage | 0.094 | 0.047 | 0.094 | 0.094 | 0.064 | 0.094 | 0.094 | 0.094 |
| Azithromycin | Inhibition of protein synthesis | 3 | 3 | 4 | 1.5 | 3 | 4 | 4 | 4 |
| Bacitracin | Interference of peptidoglycan precursor transport to cell wall | 128 | 48 | 64 | 96 | 96 | 96 | 192 | 96 |
| Clindamycin | Inhibition of protein synthesis | 0.064 | 0.125 | 0.094 | 0.125 | 0.094 | 0.125 | 0.19 | 0.125 |
| Daptomycin | Insertion into cell membrane; loss of membrane potential | 2 | 1.5 | 3 | 4 | 4 | 3 | 12 | 3 |
| Vancomycin | Inhibition of peptidoglycan cross-linkage | 1.5 | 1 | 1.5 | 1.5 | 1.5 | 1.5 | 1.5 | 1.5 |

We next wanted to examine the susceptibility of the RSV3s to two compounds commonly used to prevent oral infectious diseases: triclosan and sodium fluoride. Triclosan is a broad-spectrum antibacterial, but unlike CHX, triclosan is more compatible with other ingredients commonly used in toothpastes. Triclosan also does not stain teeth or have an unpleasant taste, as does CHX, although CHX appears to have more potent antibacterial properties (33). Triclosan was also part of a recent ban by the FDA on 19 biocidal chemicals that were deemed nonbeneficial to household wash products (34). No observable growth differences were seen when the variants were inoculated with 8 *µ*g mL^-1^ triclosan **(Figure 3A)**, but when grown with 16 μg mL^-1^ triclosan all RSV3s grew more poorly than strain UA159 **(Figure 3B)**. Only variants RSV3-3, RSV3-8 and RSV3-10 displayed final yields similar to the parental -- all others did not reach as high a final optical density. Fluoride is widely used in toothpastes and mouthwashes, and is typically added to municipal drinking water. Fluoride can not only arrest the demineralization of the tooth enamel, but at high concentrations (13 – 647 mM) it inhibits growth and metabolism of *S. mutans* (35, 36). Growth of the variants compared to the parental was moderately impaired with addition of 100 μg mL^-1^ sodium fluoride **(Figure 3C)**, but substantial growth deficiencies were observed with 300 μg mL^-1^ sodium fluoride. Collectively, these observations show that the RSV3s may display higher MICs to certain antibiotics, but they also have increased sensitivity to agents that are commonly used in oral health formulations.

**Figure 3.**
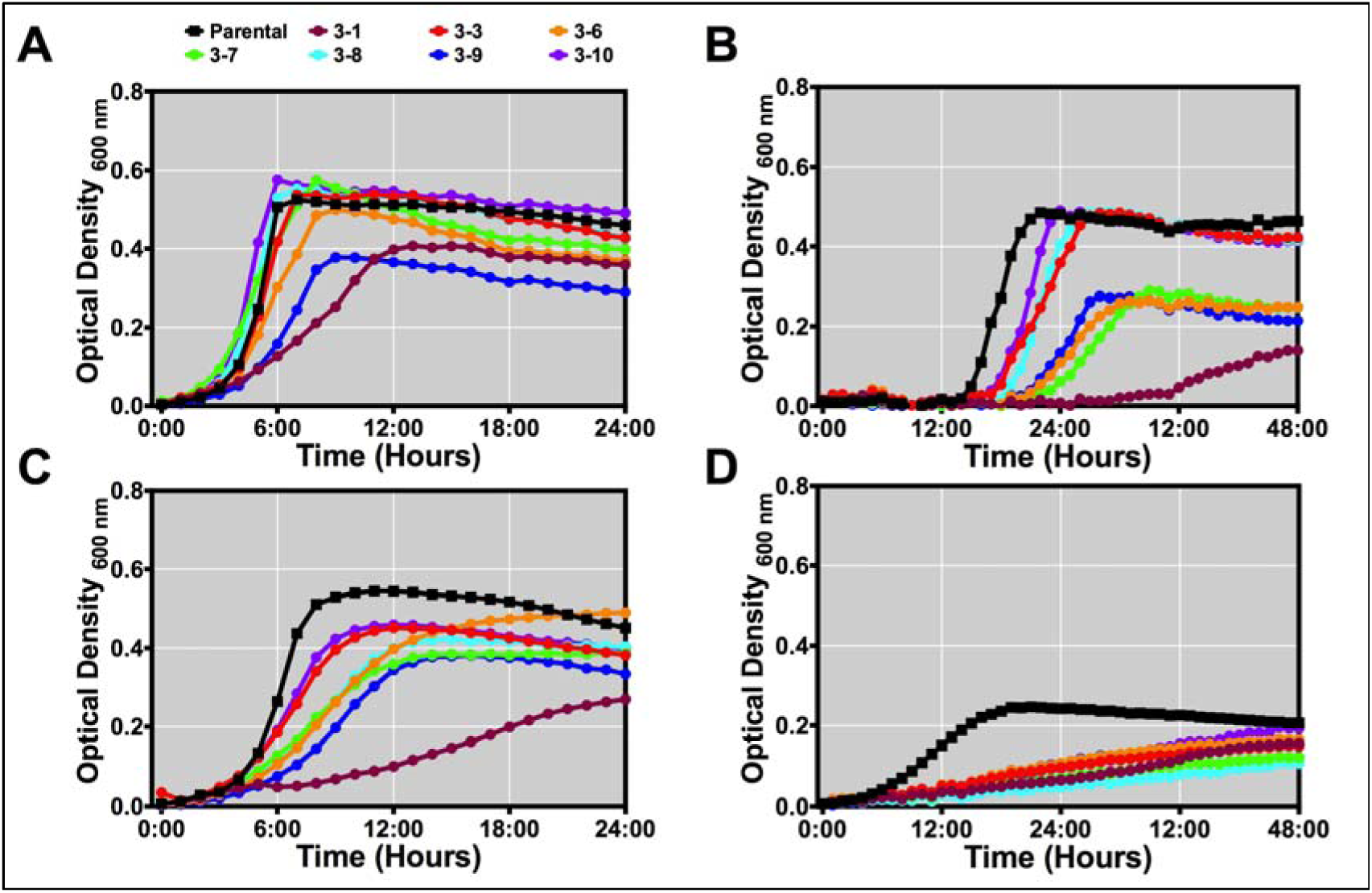
Changes in susceptibility to clinically-relevant antimicrobials. Growth of the parental UA159 strain (black line; squares) and each of the RSV3s (colored lines; circles) in either **(A)** 8 μg mL^-1^ triclosan, **(B)** 16 μg mL^-1^ triclosan, **(C)** 100 μg mL sodium fluoride and **(D)** 300 μg mL^-1^ sodium fluoride. Note differences in time scale for x-axis.

### RSV3s are Less Competitive Against Commensal Oral Streptococci

As noted above (6), there is potential for spontaneously-arising variants that show decreased susceptibility to CHX that are selected within *S. mutans* populations *in vitro*, but such isolates have not been recovered from dental plaque. One possibility is that there are selective pressures imposed by host factors and the oral microbiome that keep spontaneous CHX reduced susceptibility variants of *S. mutans* from taking hold in the human oral cavity.

To explore this idea, we investigated the antagonistic interactions of our RSV3s with species of commensal streptococci that are generally abundant and associated with dental health in humans. We measured bacteriocin production by our variants using a deferred antagonism assay in which the bacteria are stabbed on agar plates, grown overnight, and then the plates are overlaid with soft agar containing 1×10^7^ cells of a sensitive indicator strain; in this case either *S. sanguinis* SK150 or *Streptococcus gordonii* DL1. Bacteriocin production was assessed by measuring the area of growth inhibition shown by the indicator strain. When overlaid with *S. sanguinis* SK150, we observed 1.2-to 3-fold decreases in the area of growth inhibition, indicating less bacteriocin production **(Figures 4A and 4C**). A more significant difference was observed when *S. gordonii* DL1 was used as the indicator strain, as all isolates displayed ∼ 3 fold decrease compared to parental **(Figure 4B and 4D)**. We also performed a plate-based competition assay where *S. sanguinis* SK150 and our RSV3s were spotted next to each other at the same time and grown for 48 h **(Supplemental Figure 3)**. Whereas no zone of inhibition was seen in the parental when *S. sanguinis* SK150 was grown adjacent, a zone of inhibition was observed in all but two of the isolates, RSV3-1 and RSV3-8, indicating that the RSV3s did not compete as well as the parental under this condition. One source of antagonism by commensal streptococci against *S. mutans* that could account of the antagonism is hydrogen peroxide production (37, 38). To determine if the variants had marked differences in their sensitivity to H_2_O_2_ that would impact their fitness in such assays, the strains were cultured in BHI supplemented with either 0.001% or 0.003% H_2_O_2_. While differences were not observed in growth when 0.001% H_2_O_2_ was used **(Figure 4E)**, all variants displayed decreased growth rates, compared to parental, when challenged with 0.003% H_2_O_2_ **(Figure 4F)**. These data support that the spontaneous mutations that confer reduced susceptibility to CHX come at a competitive fitness cost for the variants in the form of decreased bacteriocin production and increased H_2_O_2_ susceptibility. These changes may account, at least in part, for the absence of CHX resistance among *S. mutans* isolates from human dental plaque samples. Other factor(s) not examined here may also contribute to decreased antagonism by the RS3s.

**Figure 4.**
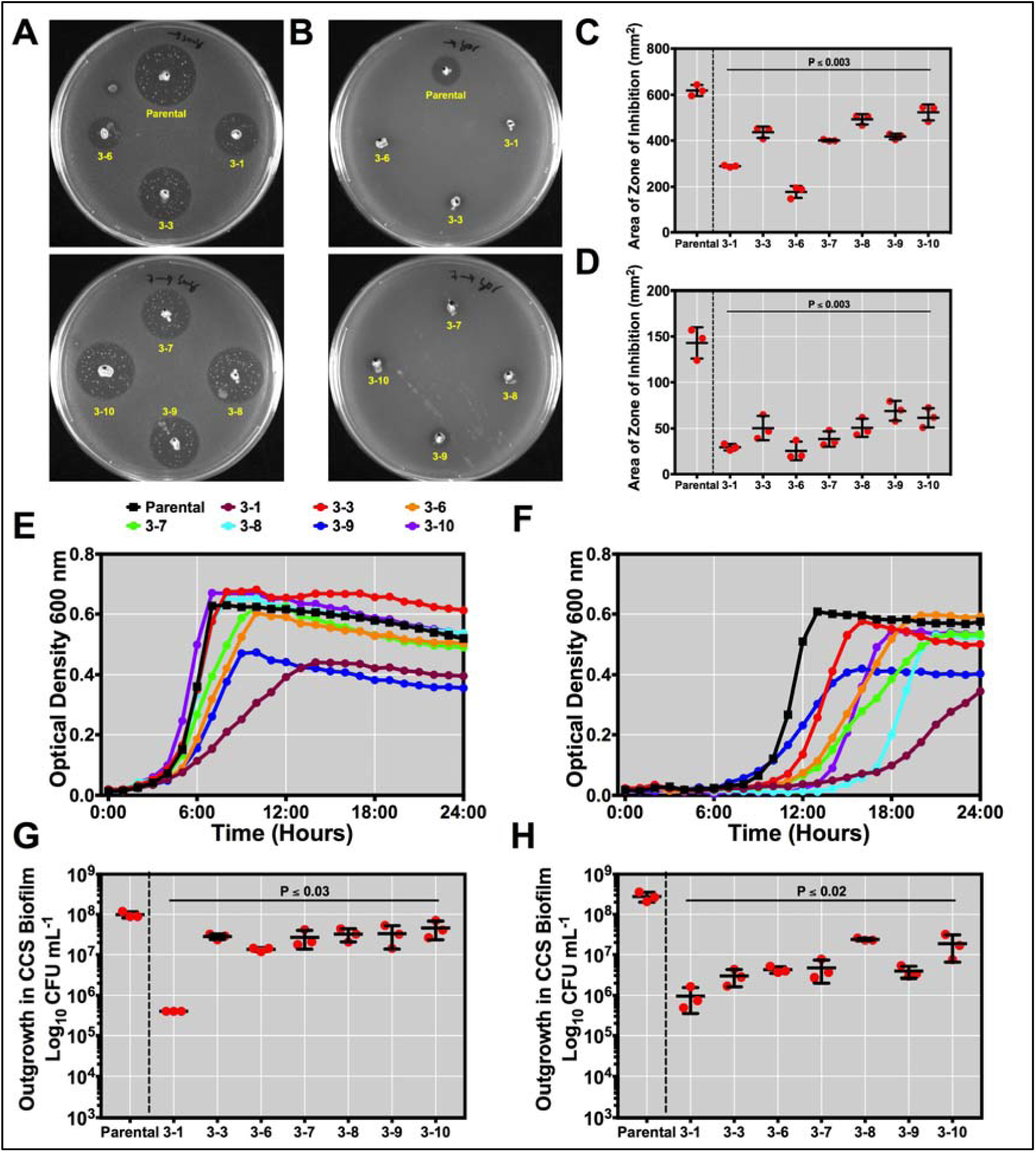
Competition of RSV3s against oral streptococci. Agar plate images of bacteriocin overlay assays displaying areas of growth inhibition against indicator strains **(A)** *S. sanguinis* SK150 and **(B)** *S. gordonii* DL1. Quantification of bacteriocin overlay assays against indicator strains **(C)** *S. sanguinis* SK150 and **(D)** *S. gordonii* DL1. Quantification was conducted using the ImageJ measuring tool and represents the average of three biological replicates from two independent experiments. All RSV3 areas of growth inhibition were statistically significant from parental (P value ≤ 0.003). Growth of the parental strain (black line; squares) and each of the RSV3s (colored lines; circles) in either **(E)** 0.001% H_2_O_2_ or **(F)** 0.003% H_2_O_2_. Outgrowth of parental and each variant in individual *ex vivo* biofilms co-inoculated with cell-containing saliva (CCS) after **(G)** 24 h and **(H)** 48 h. Outgrowth was calculated by subtracting the initial inoculation CFUs at the beginning of the experiment (t = 0 h) from the CFUs enumerated at the time point indicated. Biofilms from three biological replicates of each strain were grown. CFUs for all RSV3s were statistically significant from parental (P value ≤ 0.03 and 0.02). Each time point was replicated twice.

To further test the hypothesis that the RSV3s had diminished competitive fitness compared to the parental strain, we next examined their ability to establish and persist in an *ex vivo*, multi-species biofilm consortium. To complete this experiment, we chose to propagate the parental and each variant seperatly in independent biofilms that were co-inoculated with cell-containing saliva (CCS), collected and pooled from four healthy donors. The CCS model has been previously shown to produce a highly diverse microbial community that is predominantly composed of *Streptococcus* spp. (39–41). CFUs of each strain were determined at the time of initial inoculation (t = 0 h) and after either 24 or 48 h of incubation (t = 24 or 48 h) by plating serially-diluted cells on BHI (total cells) or mitis salivarius-bacitracin (MSB) agar supplemented with 1% tellurite and 20% sucrose (42) **(Supplemental Figures 4 and 5)**. All variants, except for RSV3-1, grew in this multi-species biofilm. We measured the outgrowth of all *S. mutans* strains, calculated by subtracting the initial inoculated CFUs at t = 0 h from the final CFUs. At t = 24 h, the RSV3 variants displayed a 2-to 7-fold reduction in outgrowth compared to the parental (mean = 4-fold) **(Figure 4G)**. The variants did not persist as well as the parental strain in the CCS biofilms at 48 h, with 12-to 94-fold lower CFUs measured (average of 55-fold) **(Figure 4H).** Total cells were also enumerated from the biofilms by plating on non-selective BHI agar and the cell count did not significantly differ between any of the samples (data not shown). The CCS assay therefore provided additional evidence that the RSV3s are impaired for growth in conditions that more closely mimic the complex communities found within the oral cavity, when compared to the parental strain. The simplest interpretation of these results is that RSV3s produce less bacteriocins, decreasing their antagonistic capability towards other streptococci present within the CCS during times of establishment, and that their ability to tolerate hydrogen peroxide over time and possibly other antagonistic factors produced by oral bacteria (37, 43) is concurrently diminished, thereby lessening their ability to persist in this model.

**Figure 5.**
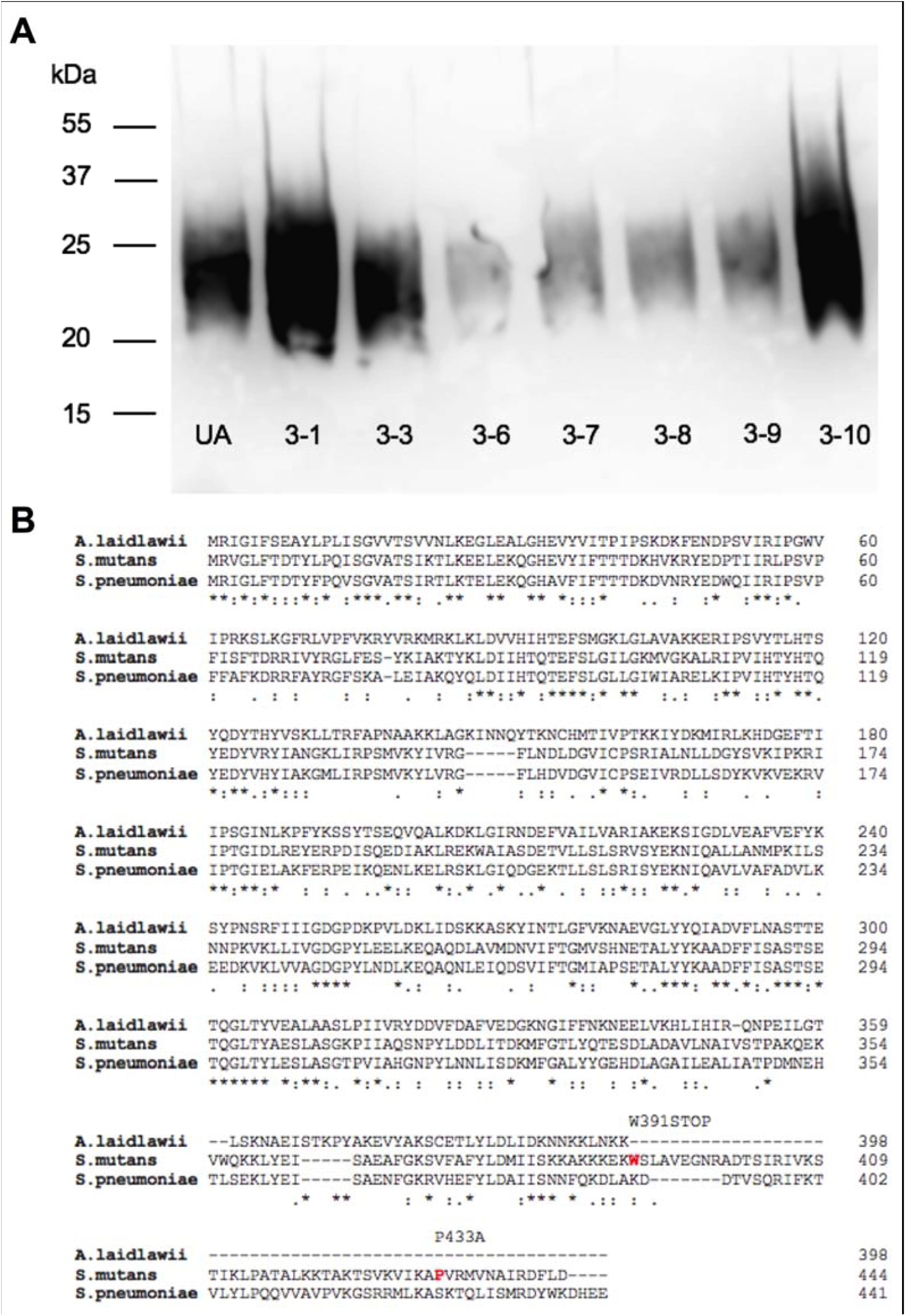
RSV3s SNPs associate with changes in LTA. **(A)** Western blot of parental and the RSV3s using a 1:5000 dilution of Gram-positive bacterial LTA monoclonal antibody. Cells were grown in BHI medium to late exponential log phase (OD_600 nm_ ∼ 0.8) prior to harvest. LTA was extracted by adding 0.5 mL of 2x sample buffer to 0.1 g of cell pellet and boiling for 1 hr. 10 μL was then added to a 4-20% TGX pre-cast protein gel. Molecular mass standards (in kDa) are shown to the left. **(B)** Amino acid (aa) alignment (Clustal Omega) between the *lafA* gene (SMU.1588c) of *A. laidlawii, S. mutans*, and *S. pneumoniae* (*D39*, spd_0961). Red bolded aa within the *S. mutans* sequence signify those residues that were found to be mutated within the RSV3s by whole genome sequence analysis, with the corresponding amino acid substitution found above. STOP = stop codon. Sequences were aligned using the web tool Clustal Omega (https://www.ebi.ac.uk/Tools/msa/clustalo/).

### Whole Genome Sequencing Reveals Mutations in RSVs

To determine what mutations are carried in each of the characterized RSV3s, whole genome sequencing was performed on extracted chromosomal DNA as described in the material and methods section. Our analysis for single nucleotide polymorphisms (SNPs) revealed four genes of interest that carried mutations in at least three or more of the sequenced RSV3s. The presence of the SNPs were confirmed by PCR and sequencing in SMU.804, SMU.1542c, SMU.1576c and SMU.1588c.

The only gene within the genome to carry SNPs in all seven of the sequenced RSV3s was SMU.1542c **(Table 3)**. SMU.1542c is homologous to *dgkB* (*yerQ*) of *Bacillus subtilis*, a soluble diacylglycerol kinase (DGK) that phosphorylates diacylglycerol (DAG) generated by the transfer of the *sn*-1-glycerol-P headgroup from phosphatidylglycerol to lipoteichoic acid (LTA) (44, 45). Intriguingly, DgkB from *B. subtilis* is essential for growth, yet four different non-synonymous SNPs were identified between our seven sequenced RSV3s that occurred over the length of the gene and were not confined to one particular region. It is not known whether the lethality of the loss of *dgkB* in *B. subtilis* is due to accumulation of toxic DAG in the membranes or from the absence of LTA (45). We suspected that the resulting non-synonymous SNPs in the apparent *S. mutans dgkB* homologue in RSV3s may result in changes to the efficiency of the enzyme, rather than complete loss of function as a recent genome-wide screen in *S. mutans* by Tn-Seq found SMU.1542c to be essential (46). To test this hypothesis, we performed a western blot on the extracted LTA from the parental strain and the RSV3s **(Figure 5A)**. Differential accumulation of LTA in the variant strains were noted. RSV3-1 and RSV3-10 appeared to accumulate more LTA and RSV3-6, RSV3-7, RSV3-8 and RSV3-9 accumulated less LTA than the parental strain. Thus, it does appear that our variants show different patterns in the amount of LTA produced that may be related to the SNPs. Further analysis will be required to determine if the LTA and lipid profiles are altered in the RSV3s, as has been reported for *E. faecium* (30).

**TABLE 3.**
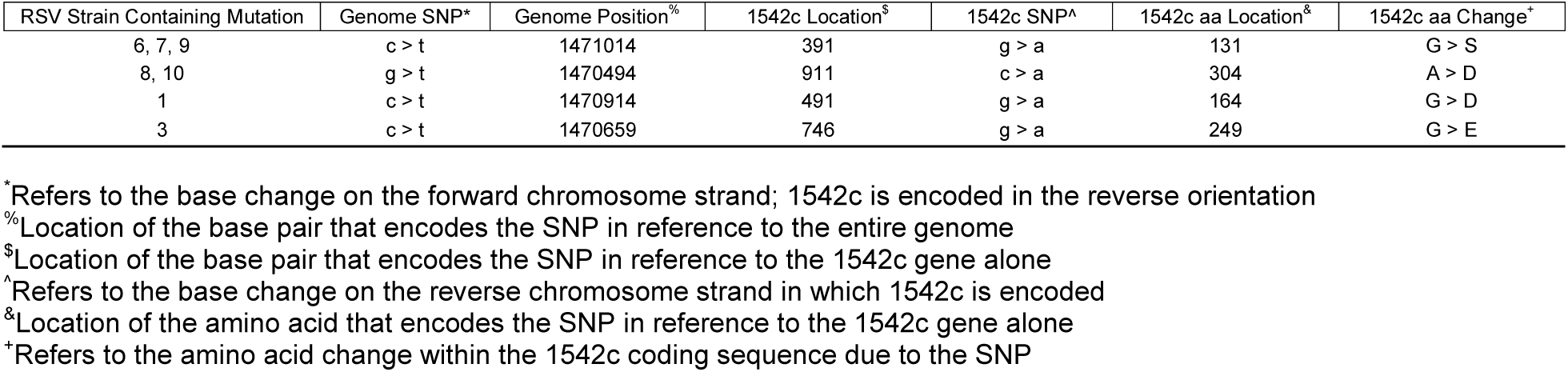
SMU.1542c Mutations in RSV3s

We analyzed the SNPs occurring within *dgkB* of our variants to determine how they might affect DgkB function within *S. mutans* by comparing to characterized Dgk enzymes, including the lipid kinase YegS from *Escherichia coli*, DgkB of *S. aureus* to which the structure has been solved (44), as well as the ten known isoforms of human DGKs **(Supplemental Figure 6)**. We noted that the amino acid (aa) substitution G131S occurring within RSV3-6, RSV3-7 and RSV3-9 arose within beta sheet 6 and was a naturally occurring substitution found within human cerimide kinase CERK, a member of the DGK superfamily (Pfam00781). The mutation causing G164D, found in RSV3-1, was a substitution occurring within DGKζ, a human type-2 DGK. Substitution D249E found in RSV3-3 occurred at the end of alpha helix 7 and is present in YegS of *E. coli*. Finally, aa substitution A304D of RSV3-8 and RSV3-10 occurred within a conserved residue of the DGKs (44) after beta sheet 17, with no known symmetry between the DGKs examined. None of the aa substitutions were within regions of the protein that impacted nucleotide (ATP/ADP)-binding regions nor along the DgkB dimerization interface. Therefore, we are unable to postulate how the identified SNPs of the RSV3s within SMU.1542c may alter the function of DgkB. However, since a majority of the SNPs lead to amino acid substitutions found in other DgkB homologs, combined with the fact that these SNPs do not lead to abberant growth defects in their respective variant as demonstrated using multiple growth tests, we assume that the DgkB enzyme in the RSV3s is still functional.

SMU.1588c is annotated to be a glycolipid synthesis enzyme, specifically a 1,2-diacylglycerol 3-glucosyltransferase that synthesizes glucose(α1-3)-DAG from uridine diphosphate glucose (UDP-glucose) and termed *lafA* in *Listeria monocytogenes* (47). It is encoded within an operon with SMU.1589c that shares homology to *lafB* of *L. monocytogenes*, responsible for the addition of a galactose moiety to form galctosyl-(α1-2)-glucosyl-(α1-3)-DAG using UDP-galactose as a substrate. In *Streptococcus agalactiae*, the LTA anchor is α1-2 linked Glu_2_-DAG and is produced by a two-enzyme system similar to L. *monocytogenes* (48–50). We encountered two different non-synonymous SNPs within the *lafA* homolog SMU.1588c: 1) aa substitution P433A in RSV3-6 and RSV3-8, and 2) W391STOP in RSV3-10, which leads to truncation of the protein at aa 391. Interestingly, the protein sequence of the well characterized *Acholeplasma laidlawii* 1,2-diacylglycerol 3-glucosyltransferase also ends at position 391 when the protein sequences are aligned **(Figure 5B)**. It is unknown what function the additional C-terminal amino acids of the *S. mutans* 1,2-diacylglycerol 3-glucosyltransferase might serve, but the additional domain(s) is present in *Streptococcus, Staphylococcu*s, *Enterococcus* and *Listeria*, but not in *Bacillus*.

Two other genes, SMU.804 and SMU.1576c, were found to have SNPs in three of the RSV3s. RSV3-7, RSV3-8 and RSV3-10 harbored SNPs in SMU.804, while RSV3-7, RSV3-8 and RSV3-9 harbored SNPs in SMU.1576c. Both genes are listed as hypothetical proteins with no known function. To determine if these two genes may be involved in a CHX reduced susceptibility phenotype that is observed in our variants, we individually replaced each gene with a non-polar kanamycin antibiotic marker to produce strains Δ804 and Δ1576c. The mutant strains did not display any changes in growth compared to the parental strain when grown in 0.50 μg mL ^-1^ or 0.75 μg mL ^-1^ CHX, nor did the loss of either gene impact the strain’s MIC to daptomycin or change antagonistic interactions with commensal streptococci as assessed with the bacteriocin overlay and plate-based competition assays **(Supplemental Figure 7)**. Together, these data suggest that while SNPs accumulated in both of these genes in three different variants, they were either not solely responsible for the phenotypes of the variants and/or the mutations did not cause complete loss of function.

Other intriguing SNPs **(Supplemental Table 1)**, such as those found in the *rcrP* gene of RSV3-1 (51) or the *rcrRPQ* peptides of RSV3-7 (52) were documented. The *rcrRPQ* operon and two peptides encoded in *rcrQ* play roles in stress tolerance and genetic competence, but SNPs in *rcrRPQ* occurred in only one of the variants sequenced. We cannot rule out, however, that these single SNPs or a combinatorial effect of the multiple SNPs in individual variants are responsible for the observed phenotypes. However, we have been unable to assign direct causation of the displayed phenotypes by any particular SNP. A future study designed to assess global transcriptomic changes in the variants compared to the parental strain may help to resolve the basis for the reduced susceptibility of growth inhibition by CHX and may provide valuable insights into mechanisms of action of the compound and ways to enhance its efficacy in oral biofilm control.

## SUMMARY

The use of antimicrobials in either mouthwashes or dentifrices in conjunction with standard mechanical removal of oral biofilms through brushing has been common practice in the dental setting since the 1970s (53). Still today, use of antimicrobials such as CHX in varnishes or even incorporation of CHX in nanoparticles is being proposed for treatments to reduce the appearance of white spot lesions in patients with fixed orthodontic appliances. Even though CHX has been familiar in the dental setting for decades, we still know very little about its mode of action against cariogenic bacteria, such as *S. mutans*, and whether or not a tolerant and/or resistant phenotypes against CHX can arise and the consequences that brings to the homeostasis of the oral microbiome. The work reported here finally begins to explore these issues and finds that after only a few rounds of passaging on CHX-containing agar can spontaneously-arising variants develop with reduced susceptibility to CHX. However, further investigation and characterization of these variants determines that while these variants do in fact have the potential to tolerate and persist at low pH values, reduced susceptibility to CHX comes at a fitness cost to the bacteria in terms of being able to compete against other commensal streptococci within oral biofilms, as well as increased susceptibility to other common oral antimicrobials such as triclosan and sodium fluoride.

While broad-spectrum antimicrobials that reduce the overall bacterial load will most likely always be found within the healthcare setting, the increase in antimicrobial resistance to common antibiotics and the realization that removal of the natural flora can have unintended consequences such as allowing pathogens into unoccupied niche(s), necessitate continuing the search for novel preventive and therapeutic interventions. The unintended finding that reduced CHX susceptibility through accumulated SNPs can lead to increased vulnerability to triclosan and sodium fluoride opens the door to future research in determining the mechanisms of action for designation of a new drug target that complements existing treatments. Together, a focus on these mechanistically-driven studies in microbial physiology and genetics will strengthen our knowledge base, provide novel insights into how we can more effectively target cariogenic bacteria, and translate to better health outcomes in the clinic to increase our quality of life.

## MATERIALS and METHODS

### Bacterial Strains and Growth Conditions

*S. mutans* wild-type strain UA159 and its derivatives **(Supplemental Table 2)** were grown in either brain heart infusion (BHI – Difco) or FMC (54) medium. The medium was supplemented with 1 mg mL^-1^ of kanamycin when needed. Unless otherwise noted, cultures were grown overnight in BHI medium with the appropriate antibiotics at 37°C in a 5% CO_2_ aerobic atmosphere. The next day, cultures were harvested by centrifugation, washed twice in 1 mL of phosphate-buffered saline (PBS), and resuspended in PBS to remove all traces of BHI. Cells were then diluted in the desired medium before beginning each experiment. For growth rate comparisons, overnight cultures were diluted 1:50 and grown to mid-exponential phase (OD_600 nm_ = 0.5–0.6). Then, cultures were re-diluted 1:100 into 0.3 mL of BHI and added to multiwell plates along with any condition to be tested (0.75 μg ml^-1^ CHX, 8 and 16 μg ml^-1^ Triclosan, 100 and 300 μg ml^-1^ NaF, and 0.001% and 0.003% H_2_O_2_). Cultures were overlaid with 50 *µ*l of sterile mineral oil to reduce the growth inhibitory effects of air on growth of *S. mutans*. The optical density at 600 nm (OD_600 nm_) was monitored at 30 min intervals for 24 h using a Bioscreen C lab system (Helsinki, Finland) at 37°C with shaking for 10 s before each reading. Three biological replicates measured in technical triplicates were measured for each experiment and the averaged growth rates of the replicates was used to calculate doubling times and final yields.

### Passaging of S. mutans on BHI Agar Plates Containing CHX

*S. mutans* wild-type strain UA159 was first grown overnight in BHI medium. The next morning, the overnight cultures were diluted 1:50 and grown to mid-exponential growth phase (OD_600 nm_ = 0.5–0.6) before being back-diluted to 1×10^6^ CFU. The 1×10^6^ CFU was plated on BHI agar plates containing either 0.1, 0.25 or 0.50 *µ*g mL^-1^ CHX (> 99.5%, Sigma-Aldrich). The colonies returned on these BHI agar plates was considered the first generation (RSV1s). 10 randomly selected colonies that appeared on the 0.50 *µ*g mL^-1^ CHX plate were then chosen for overnight inoculation to repeat the process. For the second generation, 1×10^6^ CFU of each outgrowth was plated on 10 separate BHI agar plates containing either 0.75, 1.0 or 1.25 *µ*g mL^-1^ CHX. One single colony from each of the 10 agar plates containing 1.25 *µ*g mL^-1^ CHX was then chosen for overnight inoculation to repeat the process for a third and final time. 1×10^6^ CFU of each outgrowth grown to mid-exponential growth phase was plated on 10 separate BHI agar plates containing either 1.5, 1.625 or 1.75 *µ*g mL^-1^ CHX. One randomly selected colony from each of the 10 agar plates containing 1.75 *µ*g mL^-1^ CHX was then selected to be further characterized.

### Microtiter Plate Growth Assays

CHX concentrations to be tested were first added to BHI medium with addition of DMSO only serving as a negative control. 198 *µ*l of each concentration was then pipetted across a row of a 96-well microtiter plate. Next, 2 *µ*l of either *S. mutans* wild-type strain UA159 (parental) or its derivatives to be tested grown to mid-exponential growth phase (OD_600 nm_ = 0.5–0.6) in BHI medium was used to inoculate the wells of the plate. The OD_600 nm_ of the cultures was measured after 24 h incubation at 37°C in a 5% CO_2_ aerobic atmosphere with a Synergy 2 multimode microplate reader (BioTek). The experiment was performed at least four times.

### Competitive Fitness Index Assays

To determine fitness of the RSV3s, cultures were grown overnight alongside *S. mutans* wild-type strain UA159 (parental) and UA159 harboring the chromosomally-integrated vector pMC304B that carries an antibiotic resistant kanamycin cassette which can be used for selection purposes. The next day, all strains were grown to mid-exponential growth phase (OD_600 nm_ = 0.5–0.6) after a 1:50 dilution from the overnights. A 1:1000 dilution from the mid-exponential growth phase cultures was then used to inoculate 1 mL of BHI in which either the RSV3s or parental were competed against UA159 harboring pMC304B. At this time, a small aliquot was serially diluted and plated on either BHI (total cell count) or BHI containing 1 mg mL^-1^ kanamycin (UA159 harboring pMC304B count) to determine initial cell count at the beginning of the experiment. After 24 h of growth at 37°C in a 5% CO_2_ aerobic atmosphere, cultures were again serially diluted and plated to determine final cell count. All colony forming units (CFUs) were counted after 48 h of agar plate incubation at 37°C in a 5% CO_2_ aerobic atmosphere. To determine the cell count of either the RSV3s or wild-type UA159, the UA159 harboring pMC304B count was subtracted from the total cell count. The competitive fitness index from the perspective of either the RSV3s or wild-type UA159 was calculated with the following equation: Log10 ((RSV3 cell count _final_ / UA159 cell count _final_) / (RSV3 cell count _initial_ / UA159 cell count _initial_). This experiment was repeated twice with three biological replicates for each experiment.

### Measurements of Biofilm Biomass

Selected strains were grown from overnight cultures to mid-exponential growth phase after a 1:25 dilution in CDM broth at 37°C in a 5% CO_2_ atmosphere. Cells were then diluted 1:100 into CDM medium containing 20 mM glucose and 5 mM sucrose as a carbohydrate source. 0.2 mL of this dilution was loaded into 96 well polystyrene microtiter plates and incubated in a 5% CO_2_ atmosphere at 37°C for either 24 or 48 h. After, the medium was decanted, and the plates were washed twice with 0.2 mL of sterile water to remove planktonic and loosely bound cells. The adherent bacteria were stained with 0.15 mL of 0.1% crystal violet for 15 min at room temperature. After rinsing twice with 0.2 mL of water, the bound dye was extracted from the stained biofilm using 0.2 mL of ethanol:acetone (8:2) solution. The extracted dye was then diluted 1:20 into water so that the measured values would be within a linear range of the plate reader. Biofilm formation was quantified by measuring the absorbance of the solution at OD _575 nm_ using a Synergy 2 multimode microplate reader (BioTek). This experiment was repeated twice with 3 independent biological replicates that included 4 technical replicates each.

### ETEST Antibiotic Susceptibility Testing

Selected strains were grown from overnight cultures to mid-exponential growth phase after a 1:20 dilution in BHI broth at 37°C in a 5% CO_2_ atmosphere. 10 μL from the mid-exponential culture was then added to 10 mL soft BHI agar (0.75% agarose) and overlaid on to BHI agar plates. Once solidified, ETEST strips (bioMérieux, Inc., Durham, N.C.) containing either Amoxicillin, Azithromycin, Bacitracin, Clindamycin, Daptomycin or Vancomycin when then placed in the center of the agar with sterile forceps. After 48 h of growth at 37°C in a 5% CO_2_ aerobic atmosphere, the agar plates were examined, imaged and lowest MIC recorded where no growth was visible. Two independent observers (J.K. and M.G.) interpreted the ETEST MICs. While a specific blinding protocol was not used, neither observer had knowledge of the other’s measurements prior to making their own MIC determinations. Discrepancies were resolved by consensus. Each strain was plated three times from three independent biological replicates.

### Bacteriocin Overlay Assays

Selected strains were grown from overnight cultures to mid-exponential growth phase after a 1:20 dilution in BHI broth at 37°C in a 5% CO_2_ atmosphere. Using a toothpick, the strains were stabbed into BHI agar and incubated for 24 h in a 5% CO2 atmosphere at 37°C. The following day, 10 mL soft BHI agar (0.75% agarose) was inoculated with either 10 μL *Streptococcus sanguinis* SK150 or *Streptococcus gordonii* DL1 overnight culture. The soft agar with bacterial indicator strain was overlaid onto the BHI agar plates with stabbed colonies. After 24 h, plates were imaged and the area of zones of clearance were calculated using ImageJ image analysis software. This experiment was repeated twice with three independent biological replicates in each experiment. Statistical analysis was performed using the Student’s T-Test.

### Agar Spot Competition Assays

Selected strains were grown from overnight cultures to mid-exponential growth phase after a 1:20 dilution in BHI broth at 37°C in a 5% CO_2_ atmosphere. Each strain (7 μL) was then inoculated adjacent to one another on agar plates as follows: *S. sanguinis* SK150 first and to the left, followed by *S. mutans* UA159 (parental) or RSV3 strains after the SK150 spot had dried. The plates were incubated for 48 h at 37°C, in 5% CO_2_ prior to imaging. This experiment was repeated twice with three independent biological replicates in each experiment.

### Outgrowth of Bacterial Strains in ex vivo Cell-Containing Saliva Biofilm Model

*S. mutans* UA159 (parental) and RSV3 variants from overnight cultures in BHI were diluted 1:20 into CDM medium containing 20 mM glucose and grown until the OD_600 nm_ reached 0.5. At this point, all strains were inoculated individually (n = 3) into CDM supplemented with 20 mM glucose and 5 mM sucrose using 1:1000 dilutions, along with an aliquot of cell-containing saliva (CCS) that was diluted 1:50 into CDM immediately after thawing from -80°C storage. Saliva samples for the CCS were originally collected and pooled from four healthy adult volunteers, all non-smokers that had not taken antibiotics for at least 3 months prior to collection, as described previously (IRB201500497 at the University of Florida) (55). 0.35 mL of the co-inoculated medium was loaded into an ibidi 15 *µ*-Slide 8-well glass bottom slide and incubated at 37°C in a 5% CO_2_ aerobic atmosphere for either 24 h or 48 h, while another 0.1 mL was used for serially dilution and plating on mitis salivarius-bacitracin (MSB) agar supplemented with 1% tellurite, 20% sucrose and 0.2 units mL^-1^ bacitracin to select for *S. mutans* (42) to determine initial CFU count. After 24 or 48 h of incubation, biofilms were washed twice with 1x PBS to remove loosely bound cells and then recovered by scraping off the attached cells with a pipette tip in 0.3 mL of 1x PBS and moved to a 5 mL glass tube. 0.7 mL of 1x PBS (total 1 mL) was added prior to four 30 s rounds of sonication with resting on ice in between cycles. Cell suspensions were then serially diluted and plated on MSB agar. CFUs were enumerated after the plates were incubated for 72 h at 37°C in a 5% CO_2_ aerobic atmosphere. A blank CCS control (no inoculation of *S. mutans*) was run side-by-side and plated on MSB agar plates at the beginning and end of the experiments. Between 100 and 1000 total cells were recovered, representing background levels of endogenous organisms that could grow on the selective agar. These counts were subtracted from the *S. mutans* cell counts. Outgrowth in CCS was calcuted by subtracting the initial inoculation CFU count from the final cell count and significance calculated by student’s *t*-test comparing each individual RSV3 variant to the parental. Serial dilutions of all biofilms were also plated on BHI agar to obtain a reference for total bacteria in the biofilms – the cell count of all biofilms (∼ 1×10^9^ CFUs) did not differ significantly between time points or samples. Three independent biological replicates of each strain were tested simultaneously and the experiment was performed twice.

### Construction of Bacterial Strains

Mutant strains of *S. mutans* **(Supplemental Table 3)** were created using a PCR ligation mutagenesis approach as previously described (Lau *et al.*, 2002). Briefly, 700 bp fragments both upstream and downstream of the gene of interest were amplified with designed primers and containing a BamHI restriction site in the region closest to the gene. After restriction digest, a non-polar kanamycin resistant cassette was ligated within the arms. This ligated fragment was then transformed into *S. mutans* UA159 with selection on BHI agar containing 1 mg mL^-1^ kanamycin. Successful replacement was confirmed both by size selection colony PCR and Sanger sequencing (Eurofins Genomics, Louisville, KY, USA). Restriction and DNA-modifying enzymes were obtained from New England Biolabs. PCRs were carried out with 100 ng of chromosomal DNA by using *Taq* DNA polymerase, and PCR products were purified with the QIAquick kit (QIAGEN).

### Whole Genome Sequencing and SNP Calling

For whole genome shotgun sequencing, genomic DNA were isolated from both the parental UA159 and each of the seven RSV3s using Wizard^®^ Genomic DNA Purification Kit (Promega, Madison, WI, USA) with some modifications. The total cell lysate from 10 mL of overnight cultures grown in BHI was then harvested and the supernatant containing the DNA sample was transferred to a fresh tube containing room temperature isopropanol. The supernatant was rotated at room temperature about an hour or until the thread-like strands of DNA formed a visible mass. DNA was purified in nuclease-free water after two washes in 70% ethanol. Total DNA concentration was measured using a NanoDrop™ One Microvolume UV-Vis Spectrophotometer (ThermoFisher Scientific, Waltham, MA, USA) and DNA integrity was determined by 260/280 ratio. 2-5 ng of DNA from each variant was prepared for next-generation whole genome shotgun sequencing using the Illumina Nextera-XT library preparation and indexing kit. Libraries were normalized by hand rather than with the Nextera-XT kit beads, and were pooled at a final concentration of 2 nM and sequenced on an Illumina MiSeq using the Illumina MiSeq v2 kit with paired-end sequencing and 250-bp reads. Reads were de-multiplexed by the Illumina 468 software. The pipeline a5 (58) was used to remove adapters from, quality-trim, and assemble the raw fastq files into contigs, using default settings. The quality of assembled genomes was checked using QUAST (59). The assembled contigs from each respective variant were pairwise aligned against the deposited GenBank *S. mutans* UA159 file (http://www.ncbi.nlm.nih.gov/nuccore/NC_004350.2) using the Mauve multiple genome alignment tool (version 2.4.0) (40) and SNPs unique to that variant from the parental were called through visual inspection. SNPs in genes SMU.804, SMU.1542c, SMU.1576c and SMU.1588c were confirmed via Sanger sequencing (Eurofins Genomics, Louisville, KY, USA) (Supplemental Table 3). The raw reads from all genomes we sequenced for this study are available to download from the NCBI under BioProject project accession number PRJNA515165 **(Supplemental Table 4)**.

**TABLE 4.** SMU.1588c Mutations in RSV3s

| RSV Strain Containing Mutation | Genome SNP<sup>*</sup> | Genome Location<sup>%</sup> | 1588c Location<sup>\$</sup> | 1588c SNP<sup>^</sup> | 1588c aa Location<sup>&</sup> | 1588c aa Change<sup>+</sup> |
| --- | --- | --- | --- | --- | --- | --- |
| 6, 8 | g > c | 1518523 | 1295 | c > g | 433 | P > A |
| 10 | c > t | 1518644 | 1174 | g > a | 391 | W > STOP |

<sup>\*</sup>Refers to the base change on the forward chromosome strand; 1542c is encoded in the reverse orientation <sup>%</sup>Location of the base pair that encodes the SNP in reference to the entire genome <sup>\$</sup>Location of the base pair that encodes the SNP in reference to the 1542c gene alone <sup>^</sup>Refers to the base change on the reverse chromosome strand in which 1542c is encoded <sup>&</sup>Location of the amino acid that encodes the SNP in reference to the 1542c gene alone <sup>+</sup>Refers to the amino acid change within the 1542c coding sequence due to the SNP

### Western blotting

Bacterial strains were grown in BHI at 37°C until they reached an OD_600 nm_ ∼ 0.8. Cells were harvested by centrifugation at 4000 g for 10 min, spent media was discarded, and cell pellets were stored overnight at -20°C. The following day, pellets were re-suspended in 0.5 mL 2X SDS sample buffer (100 mM Tris, pH 6.8, 4% SDS, 20% glycerol, 0.004% bromophenol blue, 10% 2-mercaptoethanol) corresponding to 0.1 g of cell pellet weight. The resuspension was then boiled for 1 h. Following, the remaining debris was sedimented by centrifugation and ten *µ*L of each sample was loaded onto a 4-20% precast TGX protein gel (BioRad). Protein samples were separated by SDS-polyacrylamide gel electrophoresis (SDS-PAGE) and then transferred to a polyvinylidene difluoride (PVDF) membrane using a Trans-Blot^®^ Turbo™ Transfer System (BioRad). LTA was detected using a Gram-positive Bacteria LTA monoclonal antibody (1:5000 primary dilution; Thermo Fisher Scientific) and a goat-anti-mouse immunoglobulin G (IgG) antibody (1:5000 dilution; SeraCare Life Sciences, USA). Western blot signals were detected using the SuperSignal™ West Pico Chemiluminescent Substrate kit (Thermo Fisher Scientific) and visualized with a FluorChem 8900 imaging system (Alpha Innotech, USA).

## ACKNOWLEDGEMENTS

We thank Dr. Robert Shields for assistance with the SNP analysis and for thoughtful discussions related to the manuscript. Research reported in this publication was supported by the National Institute of Dental and Craniofacial Research of the National Institutes of Health under Award Numbers R01 DE023339, R01 DE13239, T90 DE21990 and F32 DE028479.

---

**Table S1.**
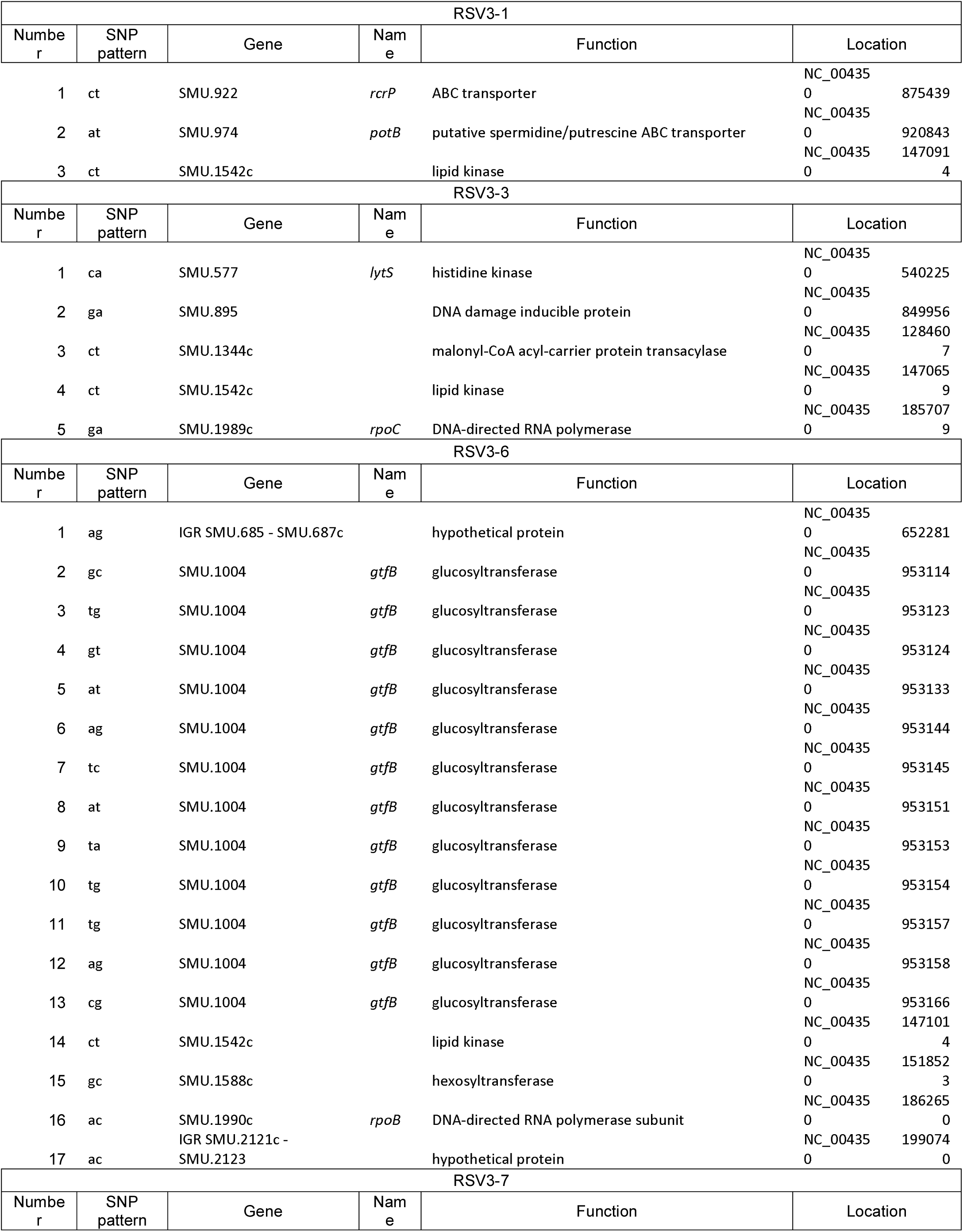
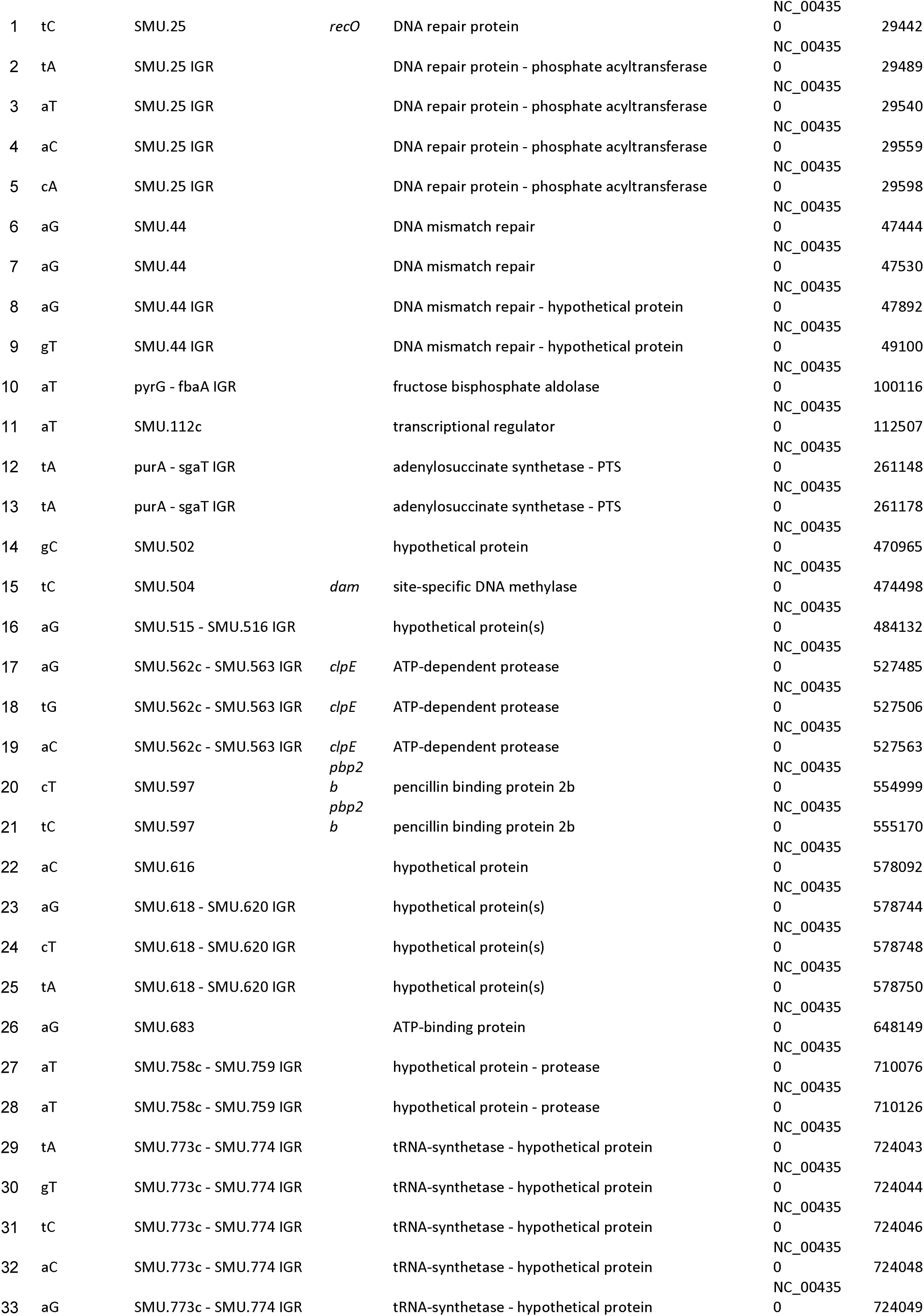
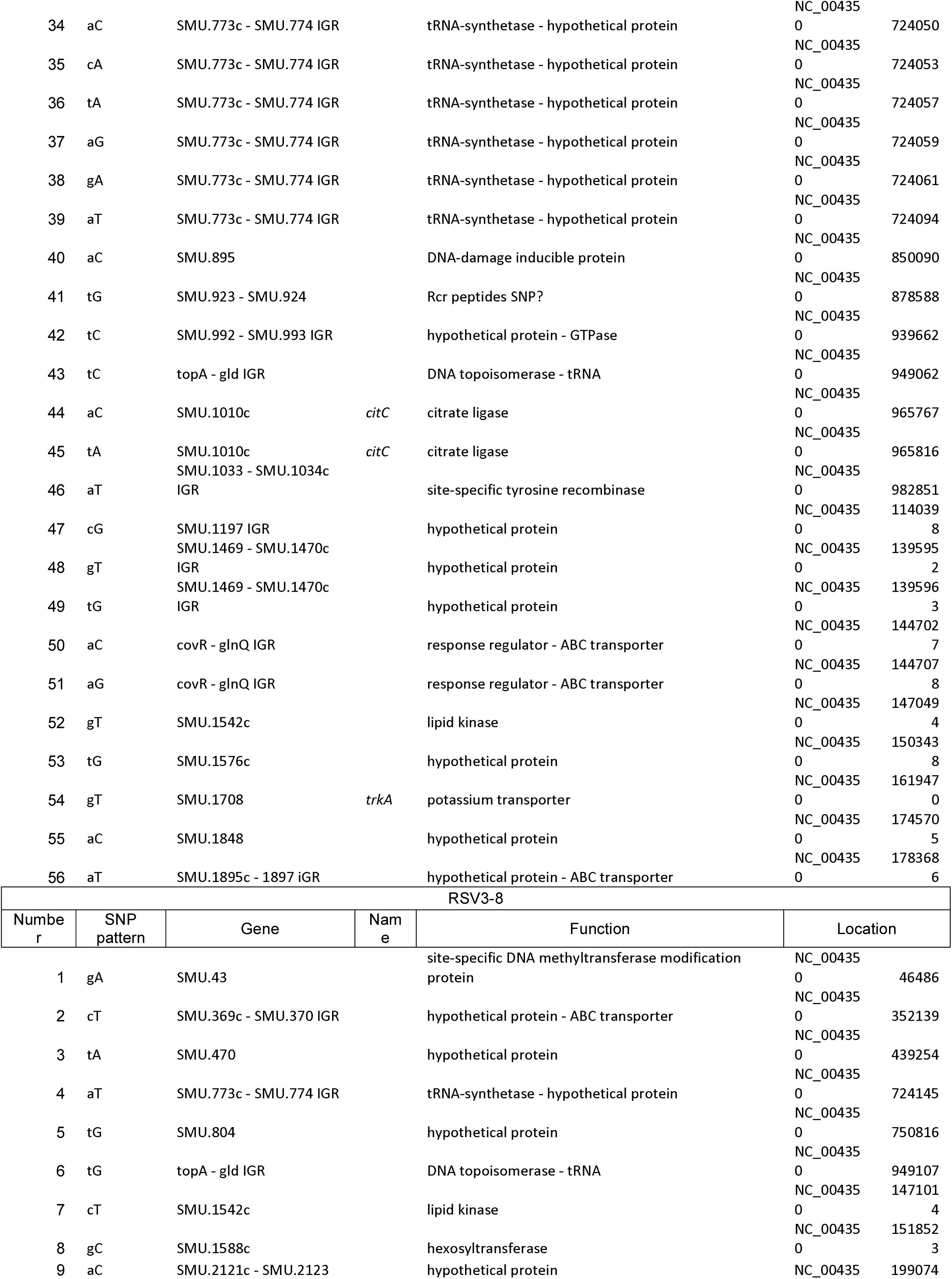
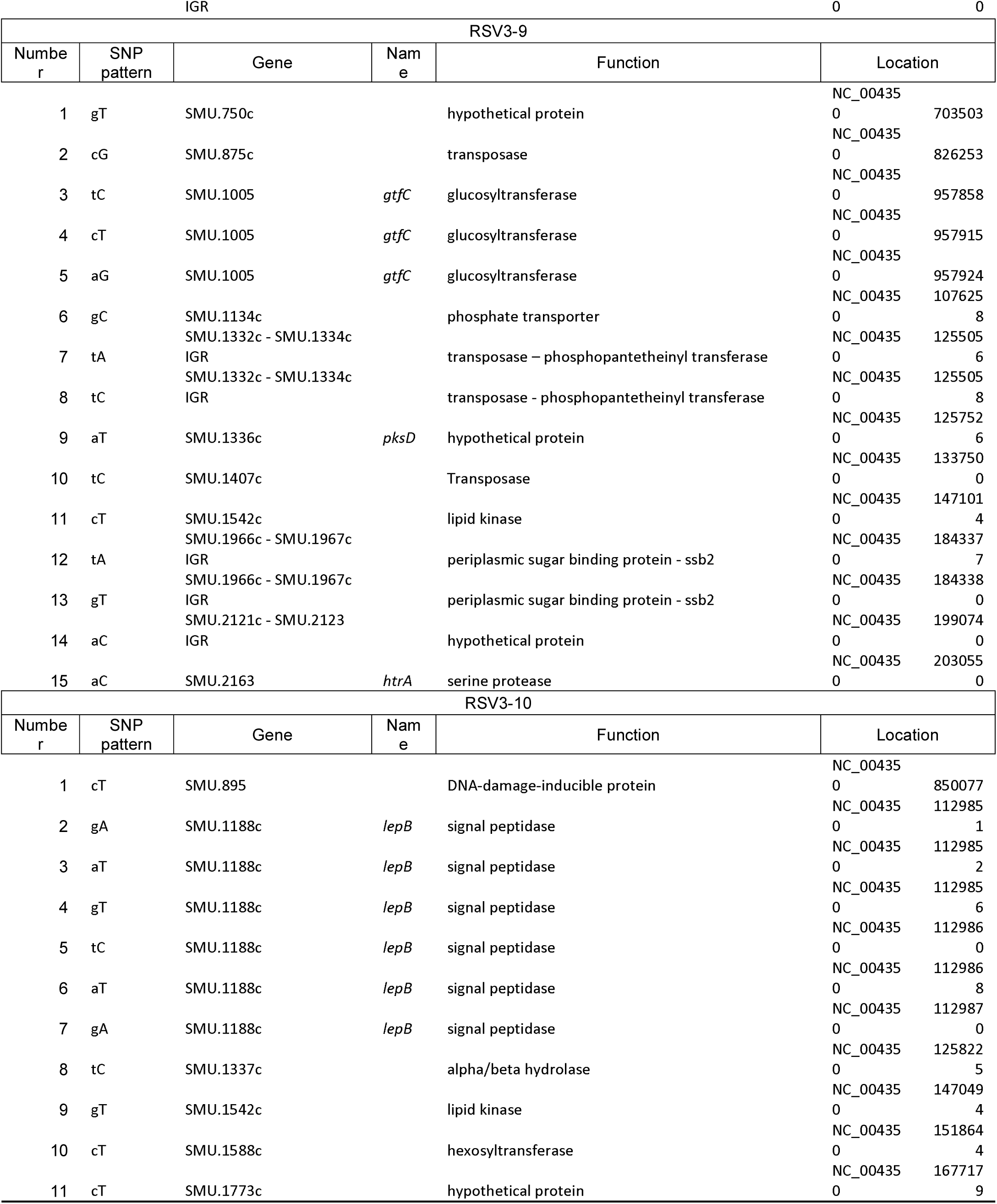
SNPs of the RSV3s.

**Table S2.** Strains Used in This Study.

| Strain | Respective Characteristics* | Reference |
| --- | --- | --- |
| *S. mutans* strains |  |  |
| UA159 | Wild-type | Lab Stock |
| RSV3-1 | Third-generation CHX passaged variant | This study |
| RSV3-3 | Third-generation CHX passaged variant | This study |
| RSV3-6 | Third-generation CHX passaged variant | This study |
| RSV3-7 | Third-generation CHX passaged variant | This study |
| RSV3-8 | Third-generation CHX passaged variant | This study |
| RSV3-9 | Third-generation CHX passaged variant | This study |
| RSV3-10 | Third-generation CHX passaged variant | This study |
| $\Delta 804$ | Non-polar Km marker replacement of SMU.804; Km<sup>R</sup> | This study |
| $\Delta 1576c$ | Non-polar Km marker replacement of SMU.1576c; Km<sup>R</sup> | This study |
| *S. gordonii* strains |  |  |
| DL1 | Wild-type | Lab Stock |
| *S. sanguinis* strains |  |  |
| SK150 | Wild-type | Lab Stock |
| *S. australis*-like strains |  |  |
| A12 | Wild-type | Lab Stock |

\* Km, Kanamycin

**Table S3.**
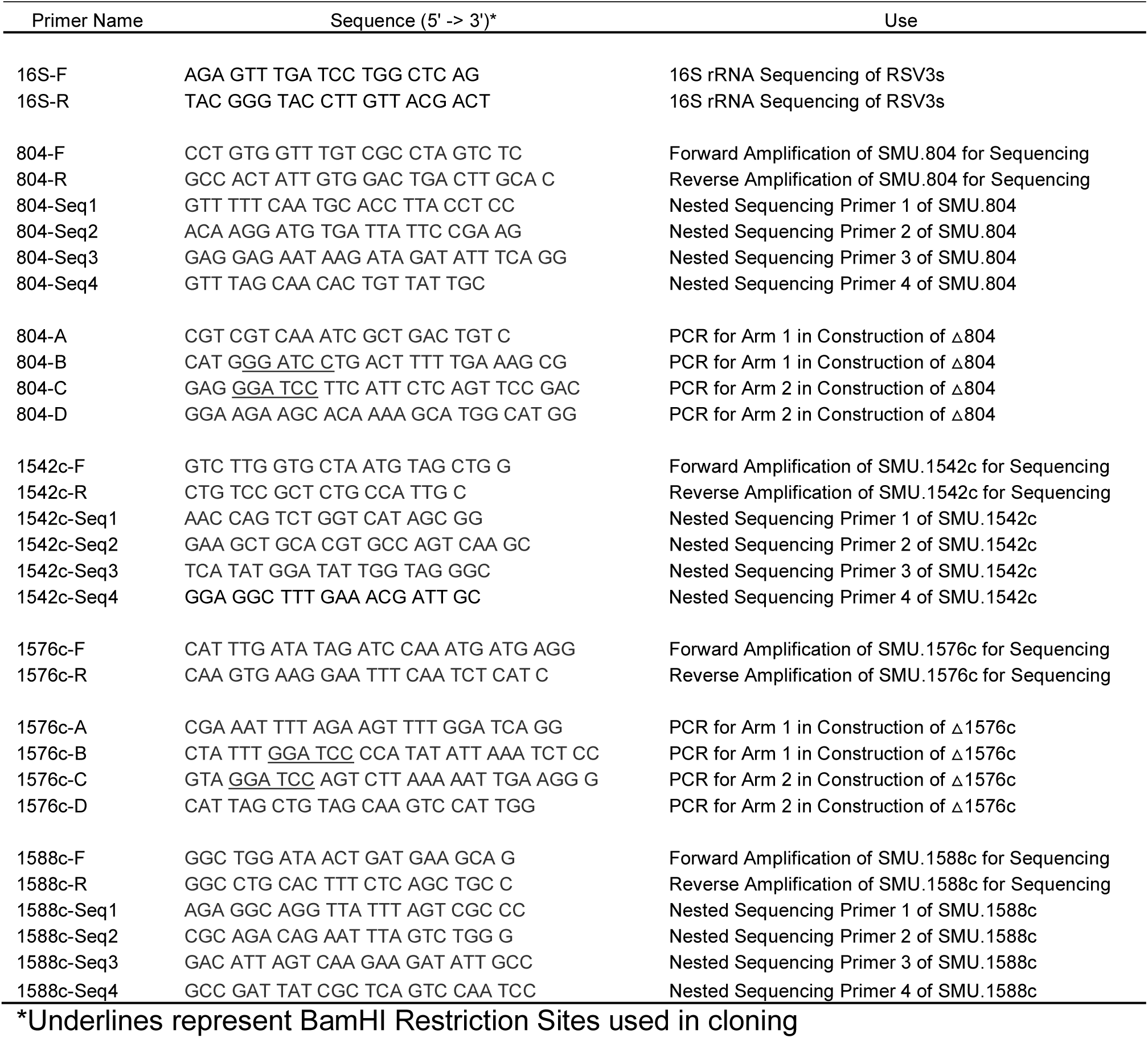
Primers Used in This Study.

**Table S4.** NCBI BioProject PRJNA515165.

| BioSample | Experiment | Sample Name | Library Name | Organism | Strain | RSV3 isolate |
| --- | --- | --- | --- | --- | --- | --- |
| SAMN10743465 | SRX5250866 | JK1 | JK1_S40 | Streptococcus mutans | UA159 | 1 |
| SAMN10743467 | SRX5250863 | JK3 | JK3_S42 | Streptococcus mutans | UA159 | 3 |
| SAMN10743470 | SRX5250857 | JK6 | JK6_S45 | Streptococcus mutans | UA159 | 6 |
| SAMN10743471 | SRX5250859 | JK7 | JK7_S46 | Streptococcus mutans | UA159 | 7 |
| SAMN10743472 | SRX5250858 | JK8 | JK8_S47 | Streptococcus mutans | UA159 | 8 |
| SAMN10743473 | SRX5250867 | JK9 | JK9_S48 | Streptococcus mutans | UA159 | 9 |
| SAMN10743474 | SRX5250866 | JK10 | JK10_S49 | Streptococcus mutans | UA159 | 10 |
| SAMN10743475 | SRX5250860 | JKW | JKW_S39 | Streptococcus mutans | UA159 | parental |

**Supplemental Figure 1.**
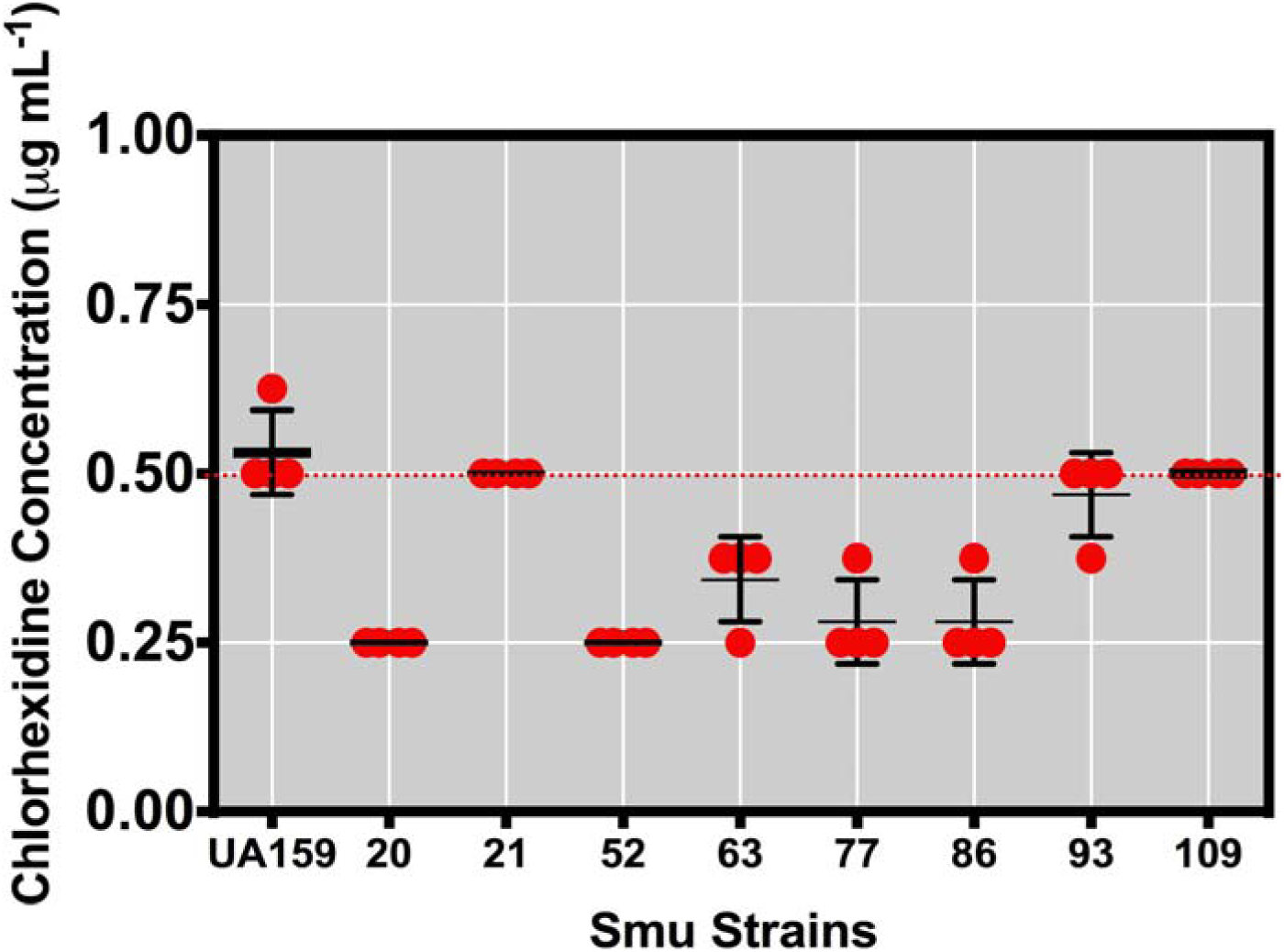
MIC Testing of Smu Isolates to CHX. Selected *Streptococcus mutans* (Smu) isolates along with the wild-type strain UA159 grown for 24 h in BHI with the designated concentrations of CHX in microtiter plates to determine minimal inhibitory concentration (MIC). Red dots denote highest CHX concentration that the strains displayed growth (OD_600 nm_ > 0.1). Four biological replicates of each isolate were tested during the experiment.

**Supplemental Figure 2.**
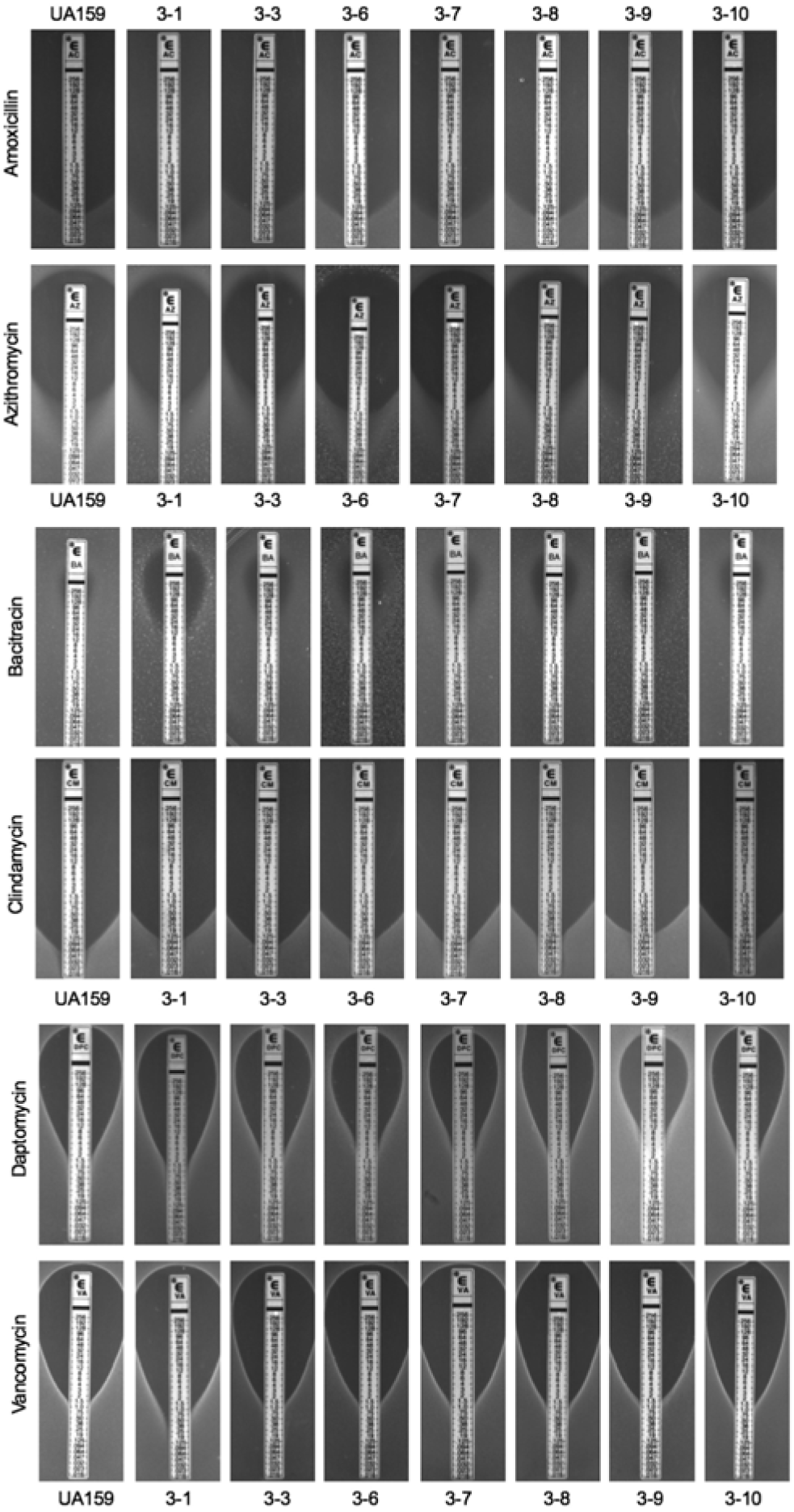
Results of MIC Testing Using ETest Strips. To determine the MIC of various antibiotics using different mechanisms of action on the parental UA159 and RSV3s, 10 μL from a mid-exponential culture of the respective strain was added to 10 mL soft BHI agar (0.75% agarose) and overlaid on to BHI agar plates. Once solidified, ETEST strips (bioMérieux, Inc., Durham, N.C.) containing either Amoxicillin, Azithromycin, Bacitracin, Clindamycin, Daptomycin or Vancomycin when then laid over the agar with sterile forceps. After 48 h of growth at 37°C in a 5% CO_2_ aerobic atmosphere, the agar plates were examined, imaged and lowest MIC recorded where no growth was visible. Each image above is one representative image from three independent biological replicates tested.

**Supplemental Figure 3.**
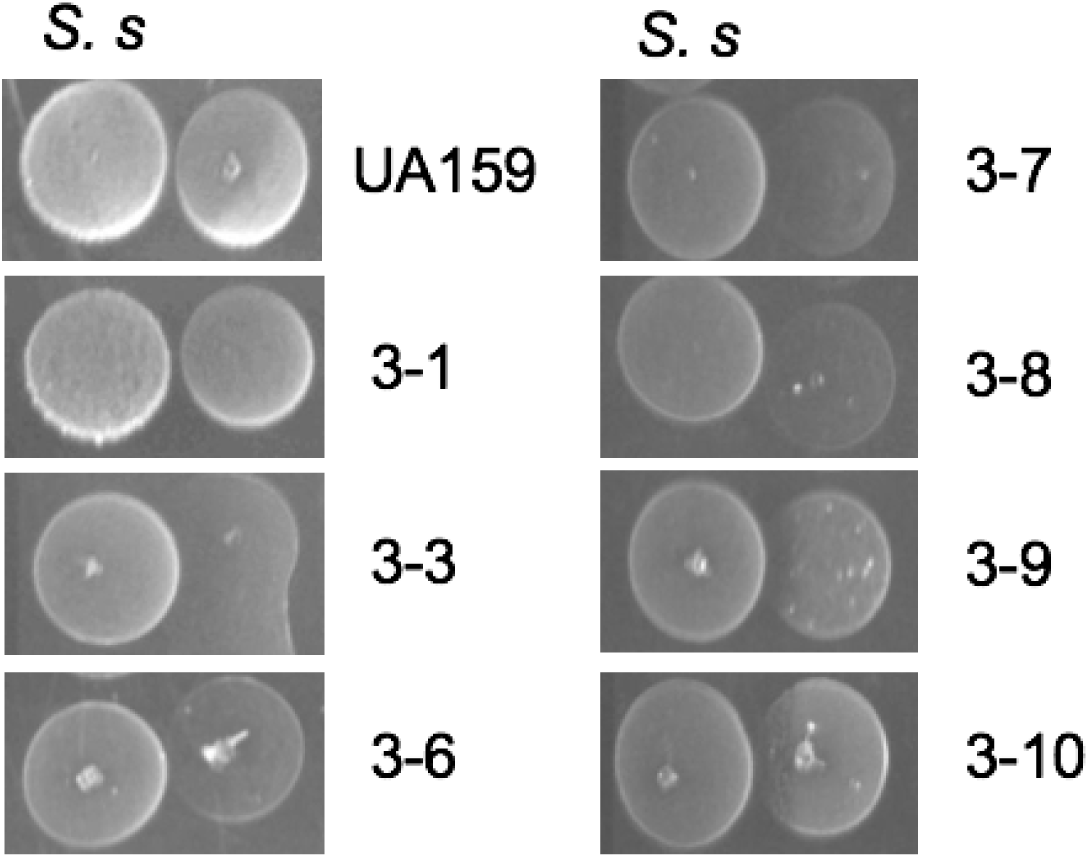
Agar Plate-based Antagonism of RSV3 Strains by *Streptococcus sanguinis* SK150. Agar plate competition assay between *S. sanguinis* SK150 (labeled S. s, left colony) and either the parental strain UA159 or the RSV3s (right colony). Colonies were spotted at the same time on BHI agar plates after growth in CDM medium to an OD_600 nm_ = 0.5. The plates were incubated for 48 h at 37°C in a 5% CO_2_ aerobic atmosphere prior to imaging.

**Supplemental Figure 4.**
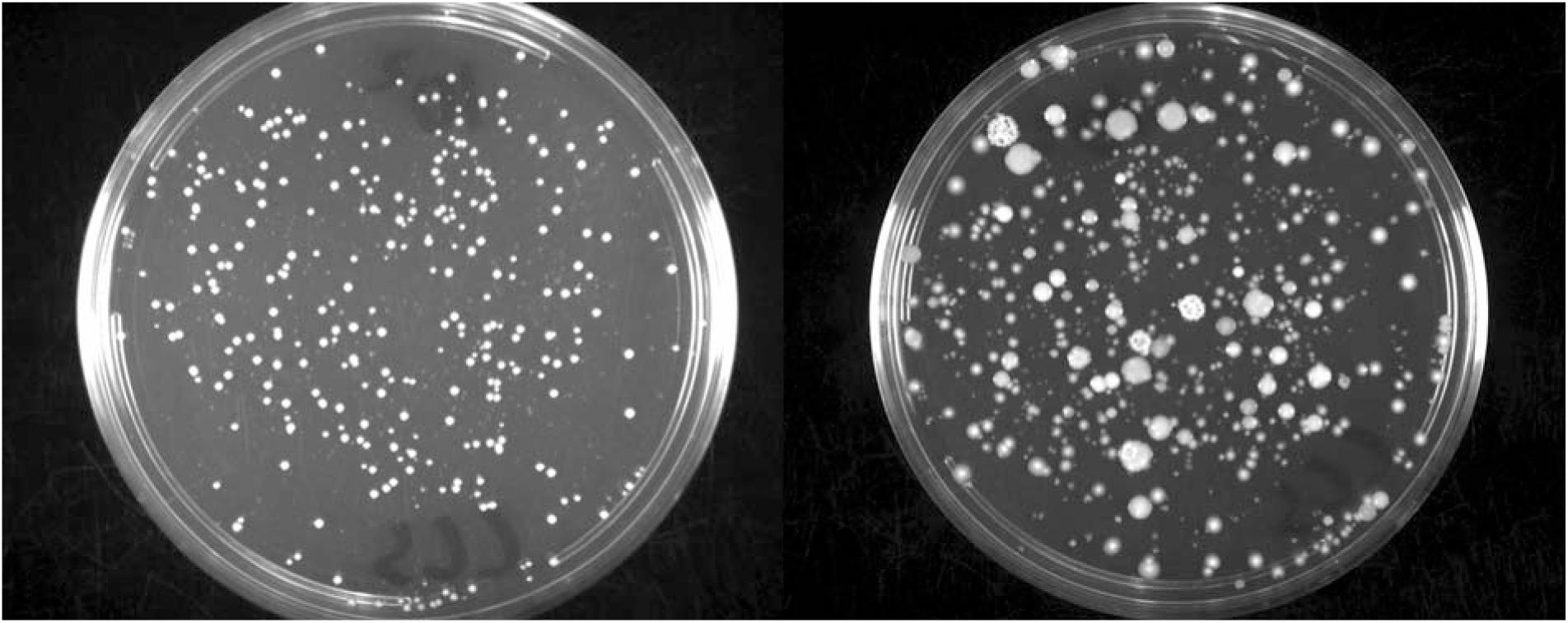
Diversity of Cell-Containing Saliva Biofilm. A cell-containing saliva control biofilm (no inoculation of parental *S. mutans* UA159 or RSV3s) was grown for 24 h in the chemically defined medium CDM supplemented with 20 mM glucose and 5 mM sucrose within an ibidi 15 *µ*-Slide 8-well glass bottom slide. After 24 h, the biofilm was washed and removed by scraping off attached cells with a pipette tip in 1x PBS. Cells were sonicated 4x 30 s prior to serial dilution and plating on BHI agar plates. Left agar plate image is the outgrowth of cells after 24 hours in incubation at 37°C in a 5% CO_2_ aerobic atmosphere, right agar plate image is 72 hours incubation.

**Supplemental Figure 5.**
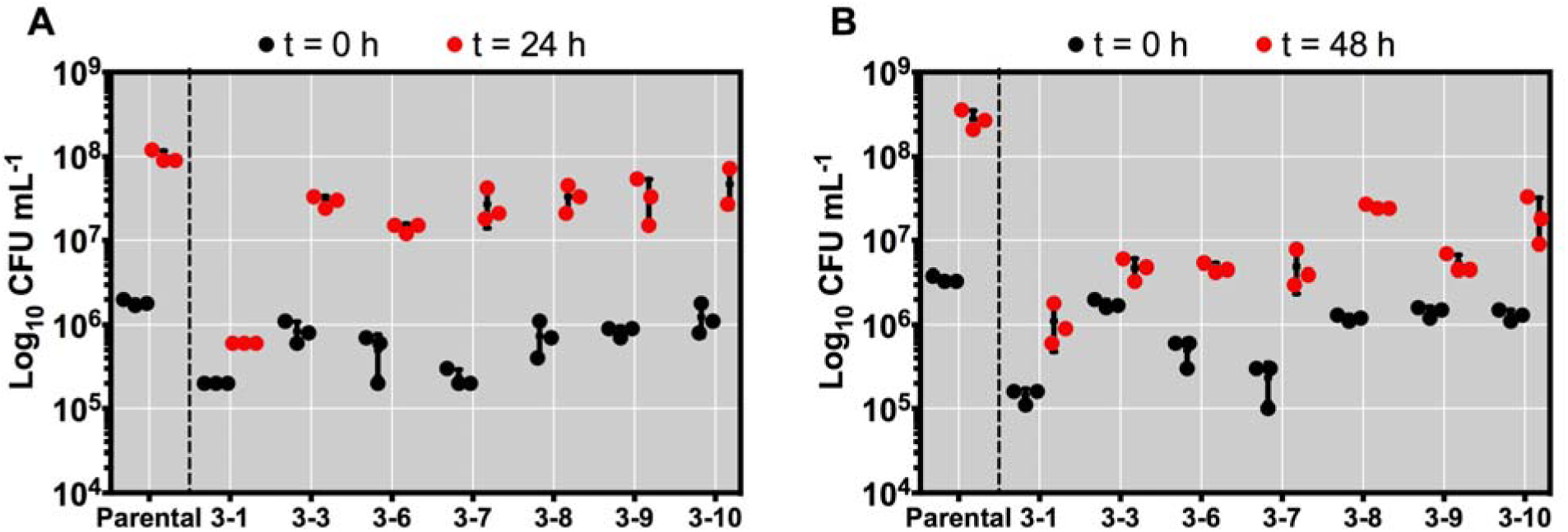
Recorded initial and final CFUs of *S. mutans* parental UA159 and RSV3s within a CCS biofilm model. *S. mutans* UA159 (parental) and RSV3 isolates were all grown in CDM medium from overnight cultures to an OD_600 nm_ = 0.5. At this point, all strains were inoculated individually into CDM supplemented with 20 mM glucose and 5 mM sucrose at a 1:1000 dilution along with an aliquot of cell-containing saliva (CCS) at a 1:50 dilution. 0.35 mL of this inoculated medium was loaded into an ibidi 15 *µ*-Slide 8-well glass bottom slide and incubated at 37°C in a 5% CO_2_ aerobic atmosphere for either (A) t = 24 h or (B) t = 48 h. Biofilms were washed to remove loosely bound cells and then removed by scraping off attached cells with a pipette tip in 1x PBS. Cells were serially diluted after sonication and plated on mitis salivarius-bacitracin agar supplemented with potassium tellurite and 20% sucrose to select for *S. mutans* only. CFUs were counted after 72 h incubation at 37°C in a 5% CO_2_.

**Supplemental Figure 6.**
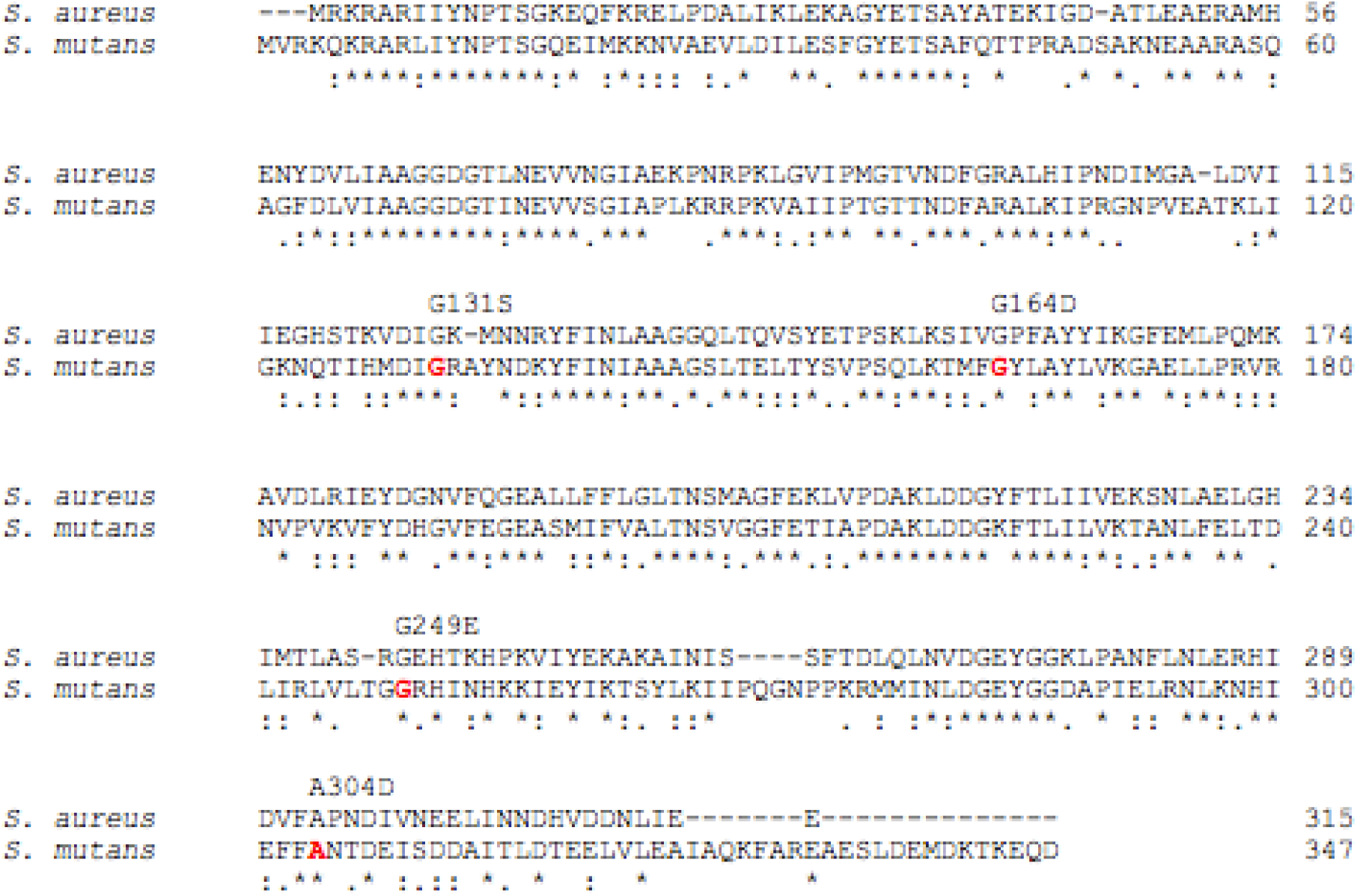
Alignment of DgkB. Alignment of the DgkB protein coding sequence between *Staphylococcus aureus* and *S. mutans* (1542c). Bolded red amino acids in the *S. mutans* sequence signify those residues that are mutated in the RSV3 variants, with the corresponding amino acid substitution found above. Sequences were aligned using the web tool Clustal Omega (https://www.ebi.ac.uk/Tools/msa/clustalo/).

**Supplemental Figure 7.**
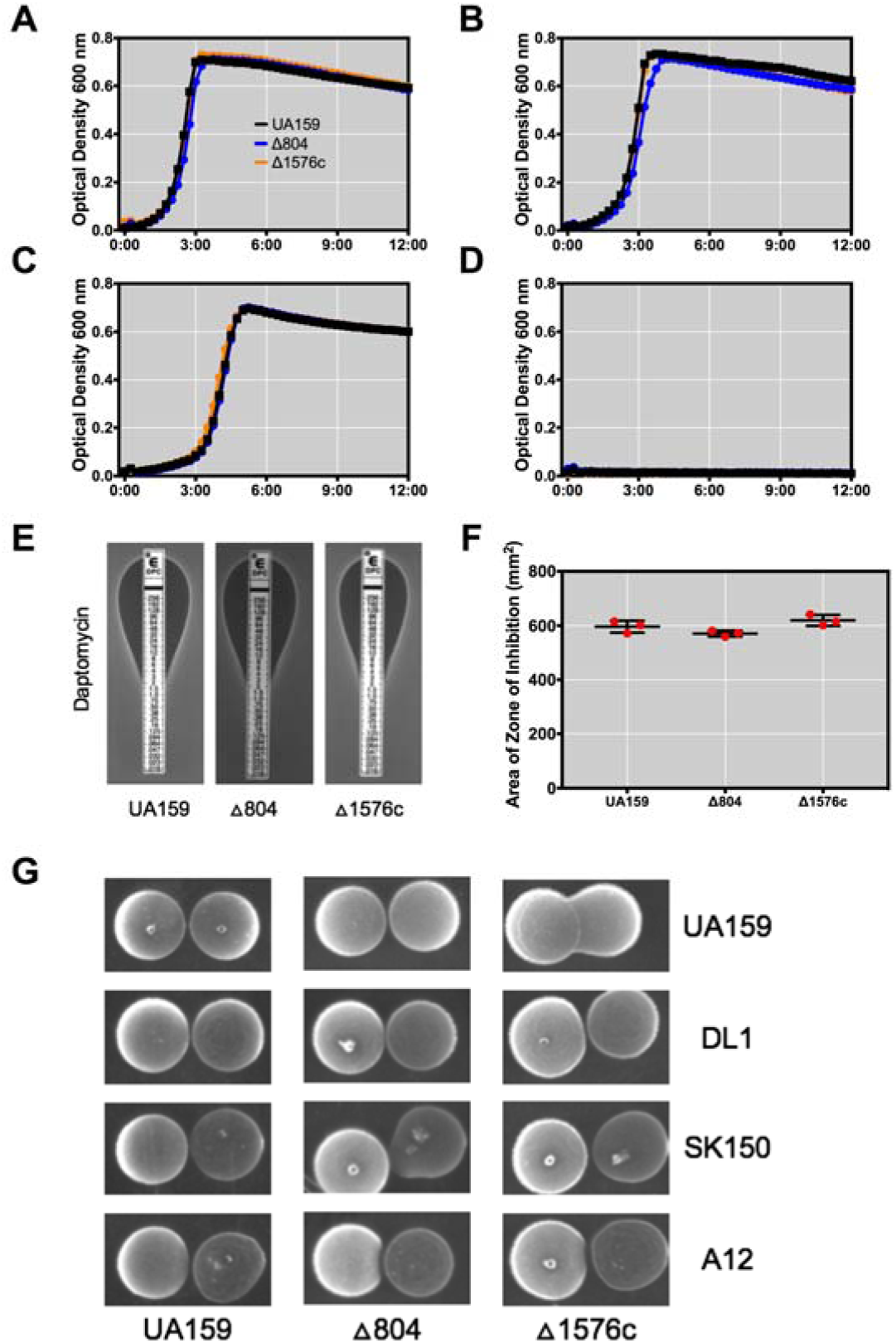
Assaying of Δ804 and Δ1576c Phenotypes. Non-polar kanamycin antibiotic marker replacement mutants of Δ804 and Δ1576c were compared to the parental UA159 strain for the following phenotypes: (A) growth in BHI, (B) growth in BHI with 0.50 μg mL ^-1^ CHX, (C) growth in BHI with 0.75 μg mL ^-1^ CHX, (D) growth in BHI with 1.00 μg mL ^-1^ CHX, (E) ETEST Strip Assay with Daptomycin, (F) bacteriocin overlay assay with *S. sanguinis* SK150 as the indicator strain, and (G) plate-based antagonism assay.

